# Linking Brain and Behavior States in Zebrafish Larvae Locomotion using Hidden Markov Models

**DOI:** 10.1101/2024.11.22.624881

**Authors:** Mattéo Dommanget-Kott, Jorge Fernandez-de-Cossio-Diaz, Monica Coraggioso, Volker Bormuth, Rémi Monasson, Georges Debrégeas, Simona Cocco

**Affiliations:** Institut de Biologie Paris-Seine (IBPS), Laboratoire Jean Perrin, Sorbonne Université, CNRS, France; Université Paris Cité, France; Laboratory of Physics of the Ecole Normale Supérieure, CNRS UMR 8023 PSL Research, Sorbonne Université, Université de Paris, France; Université Paris-Saclay, CNRS, CEA, Institut de Physique Théorique, 91191, Gif-sur-Yvette, France

**Keywords:** zebrafish, Hidden Markov Model, behavior, spontaneous neural activity, ARTR

## Abstract

Understanding how collective neuronal activity in the brain orchestrates behavior is a central question in integrative neuroscience. Addressing this question requires models that can offer a unified interpretation of multimodal data. In this study, we jointly examine video-recordings of zebrafish larvae freely exploring their environment and calcium imaging of the Anterior Rhombencephalic Turning Region (ARTR) circuit, which is known to control swimming orientation, recorded *in vivo* under tethered conditions. We show that both behavioral and neural data can be accurately modeled using a Hidden Markov Model (HMM) with three hidden states. In the context of behavior, the hidden states correspond to leftward, rightward, and forward swimming. The HMM robustly captures the key statistical features of the swimming motion, including bout-type persistence and its dependence on bath temperature, while also revealing inter-individual phenotypic variability. For neural data, the three states correspond to left- and right-lateral activation of the ARTR circuit, known to govern the selection of left vs. right reorientation, and a balanced state, which likely corresponds to the behavioral forward state. To further unify the two analysis, we exploit the generative nature of the HMM, using the neural sequences to generate synthetic trajectories whose statistical properties are similar to the behavioral data. Overall, this work demonstrates how state-space models can be used to link neuronal and behavioral data, providing insights into the mechanisms of self-generated action.

## I. INTRODUCTION

Animal behavior unfolds as a structured sequence of stereotyped motor actions, much like language. Understanding behavior thus requires identifying the vocabulary, *i*.*e*. the elementary behavioral units, and characterizing the corresponding grammar, *i*.*e*. their relative organization in time [1]. Uncovering this underlying structure is non-trivial. Over the last decade, numerous approaches have been proposed, building on the rapid development of data-driven computational methods. State-space models, in particular, appear to be well adapted, as they offer an unsupervised approach to sparse high-dimensional data into discrete states, while simultaneously unveiling their temporal structure. These include various implementations of Hidden Markov Models (HMMs) [2–5] and other statistical models [6–8].

Since behavior is driven by the brain activity, one expects the behavioral structure to be reflected in the spontaneous brain dynamics in the form of a sequence of discrete “brain states” - defined as metastable patterns of activity [9]. Neural activity can, as behavioral data, be parsed to uncover neural states and their temporal sequences [10–12] In general, however, behavioral or neuronal data are analyzed separately, as these experiments are typically conducted independently, limiting our ability to bridge the two processes. In contrast, a common modeling framework, when applied to both behavior and spontaneous neural activity, could help uncover a shared organizational structure linking self-generated neuronal dynamics and behavior.

Our model behavior is the spontaneous navigation of zebrafish larvae (see [8, 13–15] which consists of discrete swimming bouts lasting ∼100 ms and triggered at ∼1 − 2 Hz. In previous studies the categorization of bouts was carried out independently of the examination of their temporal organization. In Marques *et al*. [16], the authors used PCA-based automatic segmentation to distinguish 13 different bout types, a number that they found sufficient to encompass the entire behavioral repertoire of the animal, including hunting, escape, social behavior, etc. However, in more constrained conditions when the fish merely explore its environment [17– 23], a simple 3-state categorization is sufficient to describe their trajectories. In this case, the bouts are labeled as either forward, left-turn or right-turn based on the value of bout-induced body reorientation. The selection of these various bout types depends on sensory cues, resulting in the animal’s capacity to ascend light [17, 20] or temperature [22, 24–26] gradients.

Importantly, the neural circuit that controls the orientation of bouts has been identified as the anterior rhombencephalic turning region (ARTR), a bilaterally distributed circuit located in the anterior hindbrain. Using combined calcium imaging and motor nerve recordings, it was shown that the triggering of leftward and rightward bouts are correlated with increased activity on the corresponding side of the ARTR [18].

To characterize the behavioral and neural activities and their possible relationship, we hereafter re-analyze video recordings of freely swimming animals and ARTR recordings, performed at various water temperature, using Hidden Markov Models (HMM). First, we show that for the behavioral data, this approach provides an unbiased and therefore more consistent method of bout-type labeling compared to simple thresholding techniques as used in earlier studies. We further use the HMM inferred parameters to demonstrate and quantify inter-individual variability in exploratory kinematics. We then apply a similar 3-states HMM to the ARTR recordings performed in paralyzed tethered fish, leading to the generation of synthetic neuronal-based swimming sequences. Finally, we compare the statistical structure of these synthetic trajectories with real ones to assess the consistency of the results across both behavioral and neural data.

## II. RESULTS

### A. Data

The behavioral data used in the present article comes from a publication that examined the kinematic of free exploration in zebrafish larvae [22]. The experimental design (Fig.1a) enables recording the trajectories of multiple freely swimming larvae aged 5-7 days at temperatures of 18^*o*^C, 22^*o*^C, 26^*o*^C, 30^*o*^C, and 33^*o*^C. At each temperature, the trajectories of multiple fish are combined into a single dataset, and a set of kinematic parameters is extracted at each bout *n*, such as the angular change *δθ*_*n*_ in heading direction, the time elapsed since the previous bout and the traveled distance (see Material and Methods sec. IV A). Water temperature was found to systematically impact the statistics of navigation, leading to qualitatively different trajectories as illustrated in Figure 1b. As the temperature increases, trajectories tend to become more winding and erratic. We have also reanalyzed a second dataset of long-trajectories for 18 fish tracked individually for over two hours at 26^*o*^C, in order to assess inter-individual variability (see Material and Methods sec. IV A).

**FIG. 1.**
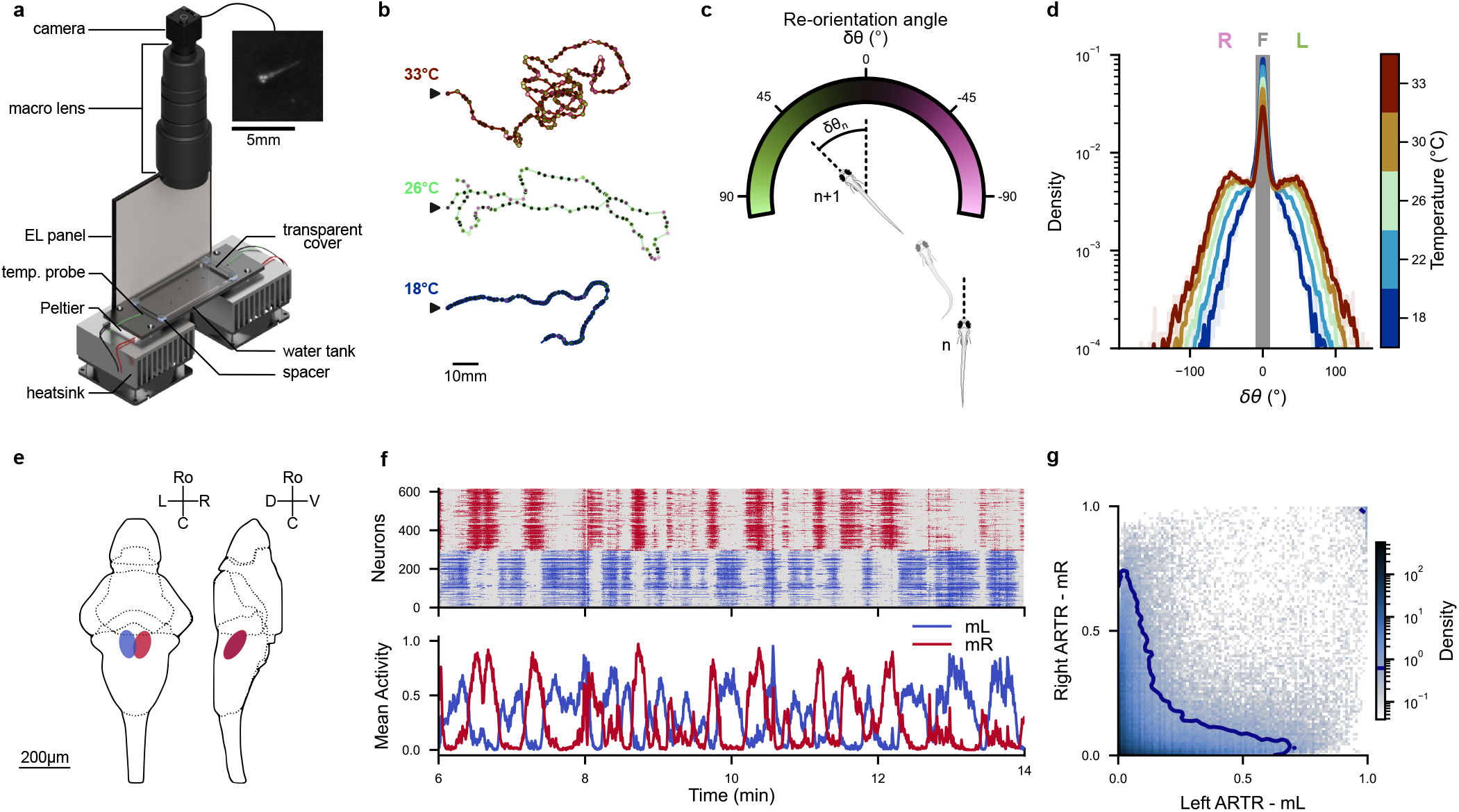
Behavioral and Neuronal Datasets: **(a)** Overview of the experimental setup: Zebrafish larvae are free to move in a tank that is kept at a desired constant temperature by a Peltier module. An imaging system records images of the fish from above at a rate of 25 frames per second. The upper right panel provides a close-up view of a larva in a raw image. Adapted from Le Goc *et al*. [22]. **(b)** Example trajectories of zebrafish larvae in 2D space at various temperatures. Each point represents a swim bout, with the color indicating the corresponding re-orientation angle defined in panel c. The trajectories’ starting points are denoted by black arrows. **(c)** Description of the convention used for the reorientation angle (*δθ*_*n*_) between two consecutive swim bouts (*n* and *n* + 1). **(d)** Distribution of re-orientation angles (*δθ*_*n*_) for each ambient temperature. The grayed-out area corresponds to the re-orientation angles classified as forward bouts by thresholds at ±10°. **(e)** Diagram of the *Anterior Rhombencephalic Turning Region* (ARTR) in larval zebrafish. Adapted from Wolf *et al*. [27]. **(f)** Example ARTR activity at 22°C. Top : Raster plot of neurons located in the left and right ARTR (blue and red respectively). Bottom : Mean activity *m*_*L*_ and *m*_*R*_ of neurons in the left and right ARTR. **(g)** Mean activities (*m*_*L*_, *m*_*R*_) of the ARTR for all recordings in the dataset. The blue contour line represents 90% of the joint distribution.

The neural data comes from another publication in which the spontaneous activity of the *Anterior Rhombencephalic Turning Region* (ARTR) [27] (Fig.1e) was recorded from 5-7 days old immobilized larvae expressing the calcium indicator GCaMP6f, using light-sheet functional imaging. Several neural recordings (3-10) for each one of the five temperatures (from 18°C to 33°C (Fig.1b)) were analyzed. The fluorescence signal of each neuron was further deconvolved to estimate an approximate spike train (see Material and Methods sec. IV B).

### B. Modeling of behavior

#### 1. Markov Models

The distribution of reorientation angles after each bout, shown in Figure 1d, appears to be trimodal, suggesting a classification of the bouts in 3 types: forward (*F*), left-turn (*L*) and right-turn (*R*). In practice, this categorization is generally carried out by thresholding the distribution of re-orientation angles. Denoting the state of swim bout n by *s*_*n*_ we have:

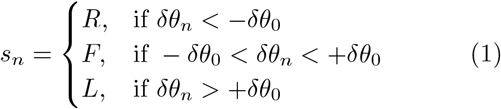

The use of the same threshold (in absolute value) to detect left and right turns relies on the hypothesis that zebrafish larvae, as a group, have no preferred direction (*a*.*k*.*a*.. non-handedness). As the exact value of *δθ*_0_ has minimal qualitative impact on the results of the Markov Chains, we adopt the same value *δθ*_0_ = 10° as in [22]; notice that *δθ*_0_ is the same across all temperatures to avoid introducing ad hoc, temperature-dependent biases. An example of the classification of states along a swimming trajectory is presented in Figure 2b.

**FIG. 2.**
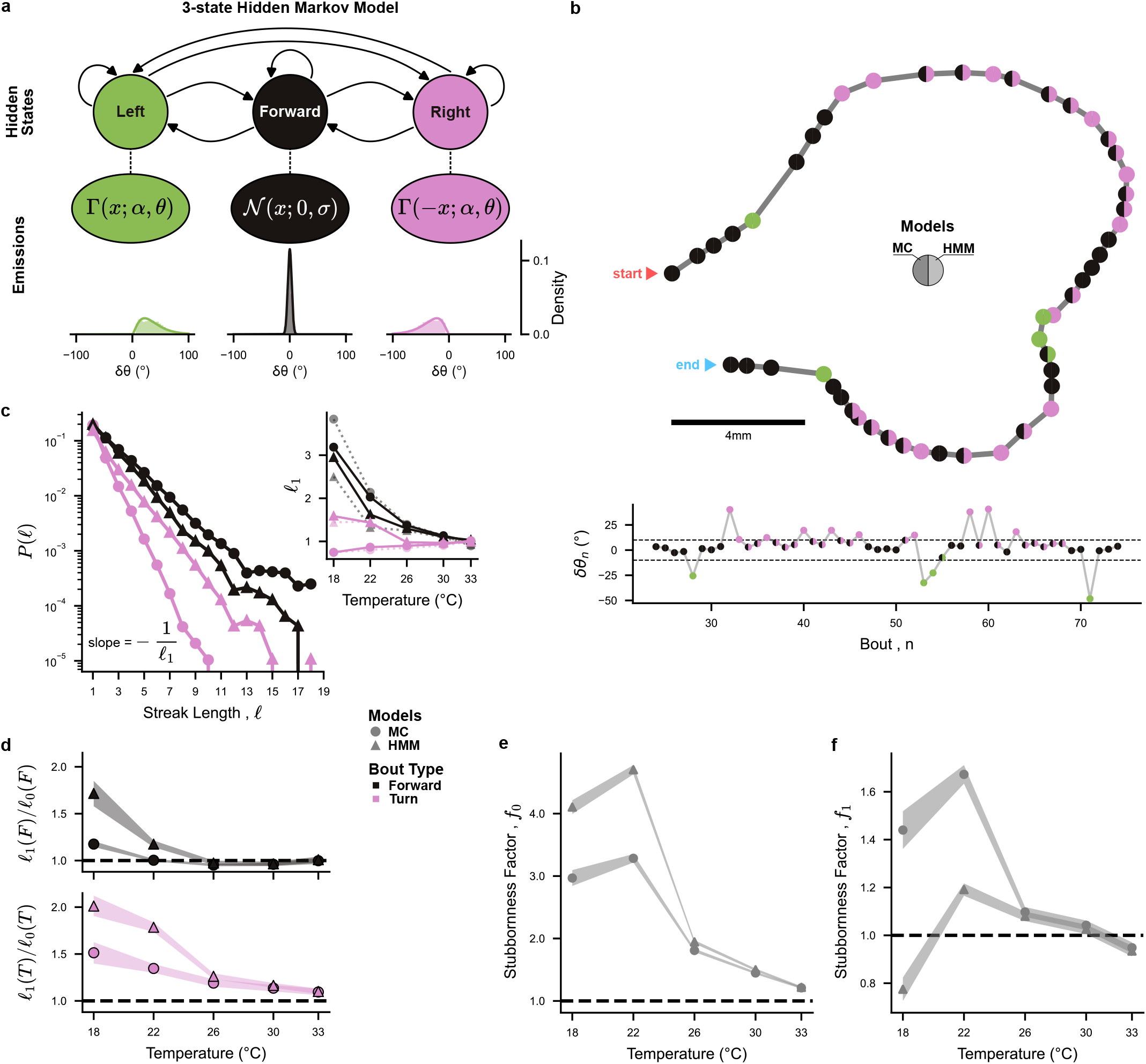
3-state Markov Chain and Hidden Markov Model - Memory effects emerge from better labeling: **(a)** Diagram illustrating the 3-state Hidden Markov Model (HMM) with emissions modeled as a normal distribution for Forward bouts, and gamma distributions for Turning bouts. Example emission distributions where taken at 26°C. **(b)** Differences in labeling between Markov Chain (MC) and HMM for an example trajectory at 22°C. Each point represents a swim bout, with the left color corresponding to the labeling according to the manual threshold used in MC, and right color indicating the HMM labeling using the Viterbi algorithm. Top: Trajectory in 2D space. Bottom: Evolution of the reorientation angle *δθ*_*n*_ for this trajectory, with the dashed lines representing the threshold *δθ*_0_ = ±10°. **(c)** Probability *P* (𝓁) of observing a streak of 𝓁 consecutive forward bouts (black) or 𝓁 consecutive turning bouts in the same direction (pink), for MC (circles) and HMM (triangles), measured from data at 22°C. Inset: Temperature dependence of the exponential decay characteristic length (𝓁_1_). Dotted line: theoretical persistence length computed from the transition matrix, 𝓁_1_(*s*) = −1*/* ln *P* (*s* → *s*). **(d)** Ratio of the observed persistence length 𝓁_1_ and the persistence expected in a no-memory null model, 𝓁_0_ vs. temperature. Forward bouts: *s* = *F*, black; turning bouts: *s* ∈ *L, R*, pink. **(e)** Temperature dependence of the *stubbornness* factor at *q* = 0 intermediary Forward bouts 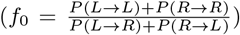. This factor is interpreted as a measurement of directional persistence during sequences of turning bouts. **(f)** Temperature dependence of the *stubbornness* factor at *q* = 1 intermediary Forward bouts 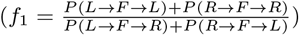. This factor is interpreted as a measurement of directional memory after one forward bout, which for a 3-state model is a second order non-Markovianity. **(e-f)** The width of the shaded curves represent the estimated error in *stubbornness* factor from aggregated fish data (see Materials and Methods IV D).

Once the bout types are identified, we define a dynamical model for the trajectories … → *s*_*n*−1_ → *s*_*n*_ → *s*_*n*+1_ → … using a three-states Markov Chain (MC). Informally, the sequence of states (associated with the 3 different bout types) is described by the probabilistic automaton in Figure S3a. In this model, after each bout *n*, a new state *s*_*n*+1_ is drawn randomly, conditioned only on *s*_*n*_ (and not on previous states). The transition probabilities between states, *P*(*s* = *s*_*n*_ → *s*′ = *s*_*n*+1_), are estimated by counting the numbers # of occurrences of the transitions *s* → *s*′ along the trajectories:

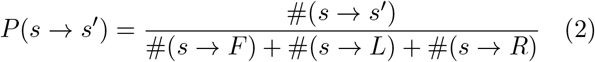

with *s, s*′ ∈ {*F, L, R*}.

The top right eigenvector of the 3 × 3 transition matrix gives access to the stationary probabilities P(*s*) of the 3 states. These probabilities are in excellent agreement with the frequencies of states estimated through direct counting (difference < 0.003 across all bout types and temperatures).

#### 2. Hidden Markov Model

We then turn to an agnostic categorization method, where states are inferred rather than *a priori* assigned. To do so, we consider a three-states Hidden Markov Model (HMM), see Figure 2a. Unlike MC, HMM makes a clear distinction between the observations (here the reorientation angles *δθ*_*n*_ treated as ‘symbols’) and the states of the system (here *s*_*n*_, which are not directly accessible from the knowledge of *δθ*_*n*_, in contradistinction with the key assumption underlying MC). The HMM is defined by:

- The transition probabilities P(s → s′) between the hidden states. We enforce the non-handedness by imposing that

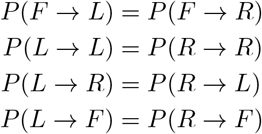

This in turn ensures that steady state bout probability is left-right symmetric (*P*(*L*) = *P*(*R*)).
- The emission probabilities, *E*(*δθ*|*s*), relate the observations δθ to the hidden states s. For the forward state, we choose normally distributed reorientation angle emission distributions, centered in zero: *E*(*δθ* |*F*) = 𝒩(*δθ*; 0, *σ*). For turn states, we use Gamma distributed reorientation angles, with a positive or negative sign according to whether the state is Left or Right: *E*(*δθ*|*L*) = Γ(+*δθ*; *α, θ*) and *E*(*δθ*|*R*) = Γ(−*δθ*; *α, θ*), constraining *α* > 1. Again, we ensured non-handedness by enforcing the same parameters for the left and right emission distribution. See Material and Methods sec. IV C for details about the validation of these emission distributions.
- A probability distribution for the initial state at the beginning of a trajectory.

We train HMM models for each dataset using the Baum-Welch algorithm, with a custom Julia [28, 29] implementation (available at https://github.com/ZebrafishHMM2023/ZebrafishHMM2023.jl/tree/bioRxiv).

### C. State classification and behavioral persistence

#### 1. Stastistics of bout states

Since the Markov Chain inferred from thresholded data (MC, Fig.S3a) and the Hidden Markov Model (HMM, Fig.2a) share the same internal behavioral states, we can directly compare these two models and thus examine the impact of the labeling methods.

As illustrated with an example trajectory at 22°C in Figure 2b, MC and HMM labeling can differ significantly. MC-inferred sequences often exhibit multiple alternations between Forwards and Turns when the bouts reorientation angles are near the threshold, while for the same sequence, the HMM tends to consistently label these bouts as Turns. These differences result in a reclassification of approximately 60% of Forward bouts into Turning bouts at 22°C (Fig.S3e).

The HMM yields a relatively modest dependence of bout-type usage on temperature (see Fig.S3b). In contrast, the hard-threshold classification method used in MC lead to a systematic and pronounced increase in the fraction of turning bouts with rising temperature. This strong temperature dependence, previously reported in Le Goc *et al*. [22], may have thus been overestimated, as it partly reflects the ad-hoc assumption of a fixed (temperature-independent) threshold *δθ*_0_. Conversely, the HMM approach infers a gradual widening of the forward bouts angular distribution with increasing temperature that effectively corresponds to an increase in the angular threshold (see Fig.S2c-e).

#### 2. Bout streaks and persistence

We further assessed how bout-type persistence, defined as the tendency to execute similar bouts in succession, depends on the chosen classification model. We start by describing trajectories as a series of streaks of similar bouts (forward, leftward or rightward), and then characterize the streak length distribution. For all bout types and models, the probability of observing a streak of 𝓁 consecutive bouts of the same type decays exponentially, 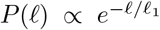, with 𝓁_1_ defining the characteristic streak length (Fig.2c). For turning bouts, we found 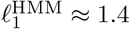 bouts while 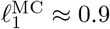 bouts at 22°C. Compared to MC, HMM-based labeling thus yield much longer turning streaks. In contrast, we find no significant difference in characteristic forward-streak length between HMM and MC. As temperature increases, we observe for both models that the characteristic streak length decreases (particularly for forward bouts, see Fig.1b).

Within the Markov or Hidden Markov Model frameworks, the average length 𝓁_1_(*s*) of a streak of bouts of type *s* is related to the probability *P*(*s* → *s*) of remaining in the same state through the simple relation 𝓁_1_(*s*) = −1/ ln *P*(*s* → *s*). To distinguish the effects on bout-type persistence due to the presence of memory from the mere consequences of single-state frequencies, we introduce a null model, in which the transition probabilities are simply given by these frequencies, *i*.*e. P*(*s* → *s*′) = P(*s*′). In this null model without any memory, the average length of type-s bouts is simply 𝓁_0_(*s*) = −1/ ln *P*(*s*). The ratio 𝓁_1_(*s*)/𝓁_0_(*s*) is an estimator of the (relative) contribution of behavioral memory to bout-type persistence.

Results for this memory-induced persistence are shown in Figure 2d for the Markov (MC) and Hidden Markov (HMM) Models. The MC and HMM methods yield comparable outcomes for turning bouts at low temperature. However, HMM-based analysis further reveals a persistence for forward bouts at lower temperatures (Fig.2d), while this effect is absent in the MC model. Here again, this absence of forward persistence, previously reported in Karpenko *et al*. [20], is likely due to the mis-labeling associated with the hard-threshold method. Interestingly, such persistence effects vanish at higher temperatures, where the transition matrix becomes uniform (Fig.S3c,d), and all bouts become equiprobable (*P*(*F*) ≈ *P*(*L*) ≈ *P*(*R*)). One thus expect more erratic trajectories at higher temperatures, which is consistent with our observations (see Fig.1b).

#### 3. Consistency of the MC and HMM descriptions of behavior

Taken together, the results above suggest that the Hidden Markov Model better captures persistence in reorientation by labeling bouts with small reorientation angles based on context. This leads to a more flexible and thus stable classification than the hard-thresholding method. However, given the absence of a ground truth, it remains unclear whether the labeling produced by the Hidden Markov Models is more accurate than the one produced by the standard threshold-based approaches.

One way to address this question is to examine to what extent each of these methods are self-consistent, guarantees that the inferred labeled sequences are truly markovian such that the bout type at a given time only depends on the type of the preceding bout. It has been previously noted that the hard-thresholding methods lead to significant non-markovianity. In particular, in a transition *T*_1_ → *F* → *T*_2_ with *T*_1_, *T*_2_ ∈ {*L, R*}, the two turning bouts tend to have the same orientation (*T*_1_ = *T*_2_). This means that the memory of orientation *T*_1_ is maintained during the forward bout, in violation of the Markovian assumption. This observation lead to propose a 4-state Markov system comprising two independent Markov chains, independently controlling the forward-turn bout transitions, and directional left-right bout transitions (see Fig.S4b for a diagram of this 4-state model) [20, 22].

Given that our 3-state Hidden Markov Model (HMM) re-labels numerous Forward bouts as Turn bouts, we ask whether this new classification might alleviate this non-Markovianity issue, such that the ad hoc 4-state model might no longer be needed. We thus propose a new test of Markovian violation specifically designed for our use case, that we apply to both the HMM and MC models.

We introduce the *stubbornness factor f*_*q*_ to empirically assess the tendency of larvae to retain their orientation after a sequence of q intermediary forward bouts (Fig.S4b, Materials and Methods sec. IV D):

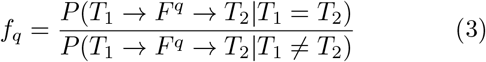

with *T*_1_, *T*_2_ ∈ {*L, R*} and 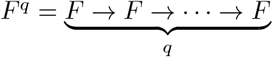.

Owing to the loss of orientational memory after a forward bout, a non-handed 3-state Markovian model should have *f*_*q*_ = 1 for *q* ≥ 1 (Materials and Methods sec. IV E). On the other hand, *f*_*q*=0_ is a measurement of directional persistence during uninterrupted sequences of turning bouts.

We found that most of the memory effects captured by the HMM occur at q = 0, and that the *stubbornness* reaches *f*_*q*_ ≈ 1 for *q* ≥ 1, suggesting that the HMM-inferred bout sequences are quasi-markovian. In comparison, and for lower temperatures, the thresholded MC classification displays lower persistence at q = 0 but higher *stubbornness* at *q* = 1 as seen on Figure 2e-f (and less significantly at *q* = 2, see Fig.S4d). This suggests that the thresholded labeling leads to Markov violation primarily due to the mislabeling of turn bouts as forward bouts during turning streaks, as anticipated in the previous section and illustrated on Figure 2b. As this *stubbornness* is mostly significant at *q* = 1, we expect that most mislabelings are one-off errors.

In summary, previous works using a ad hoc threshold to classify bouts had dismissed 3-states Markov models because the resulting sequences were non-markovian. We found that by using an unsupervised method to simultaneously label the data and infer a Markov Model, we could unveiled previously underestimated memory effects in zebrafish reorientation statistics. Our results suggest that the apparent non-markovianity reported in previous studies was mainly caused by the mislabeling of turning bouts as forward bouts during sequences of consecutive turns. The HMM seems to be a clear improvement, identifying quasi-Markovian 3-state sequences and providing a more robust representation of the swimming dynamics.

### D. Behavioral phenotyping from long individual fish trajectories

As HMM provides an unbiased quantification of the behavior, we now ask whether the approach is accurate enough to detect behavioral differences between specimen (inter-individual variability) and whether it can enable the unambiguous identification of each animal.

In the preceding sections, the dataset used to infer the models comprised trajectories from multiple fish, as the different individuals swimming together during a given assay could not be distinguished. To address the question of individuality, we used additional experiments reported in Le Goc *et al*. [22], in which individual fish were tracked at 26°C (see Materials and Methods IV A). A total of 18 fish were recorded for over 2 hours.

We first split the 2h-long recorded sequence of each individual fish into smaller periods (chunks) of ≈12 minutes each, and trained an HMM on each of these chunks (see diagram in Figure 3a-b). For each fish, the parameters of these HMMs exhibit significant variability (as shown by the vertical error bars in Figure 3c). This variability between the different chunks reflects both intra-individual (temporal) variability and, to a lesser extent, inference uncertainty due to the limited sampling of the HMM (see Fig.S5). We then also trained a single HMM on the entire dataset of a single fish (the “global” HMM). Figure 3c compares selected parameters of the global HMM for each fish, against the average parameters over several HMMs trained on the chunk trajectories (see Fig.S5 for all parameters). There is a clear trend between the global HMM and the average behavior of the chunk HMMs. Therefore, although a fish exhibits variability during a long sequence of bouts, the variability between distinct fish is larger.

**FIG. 3.**
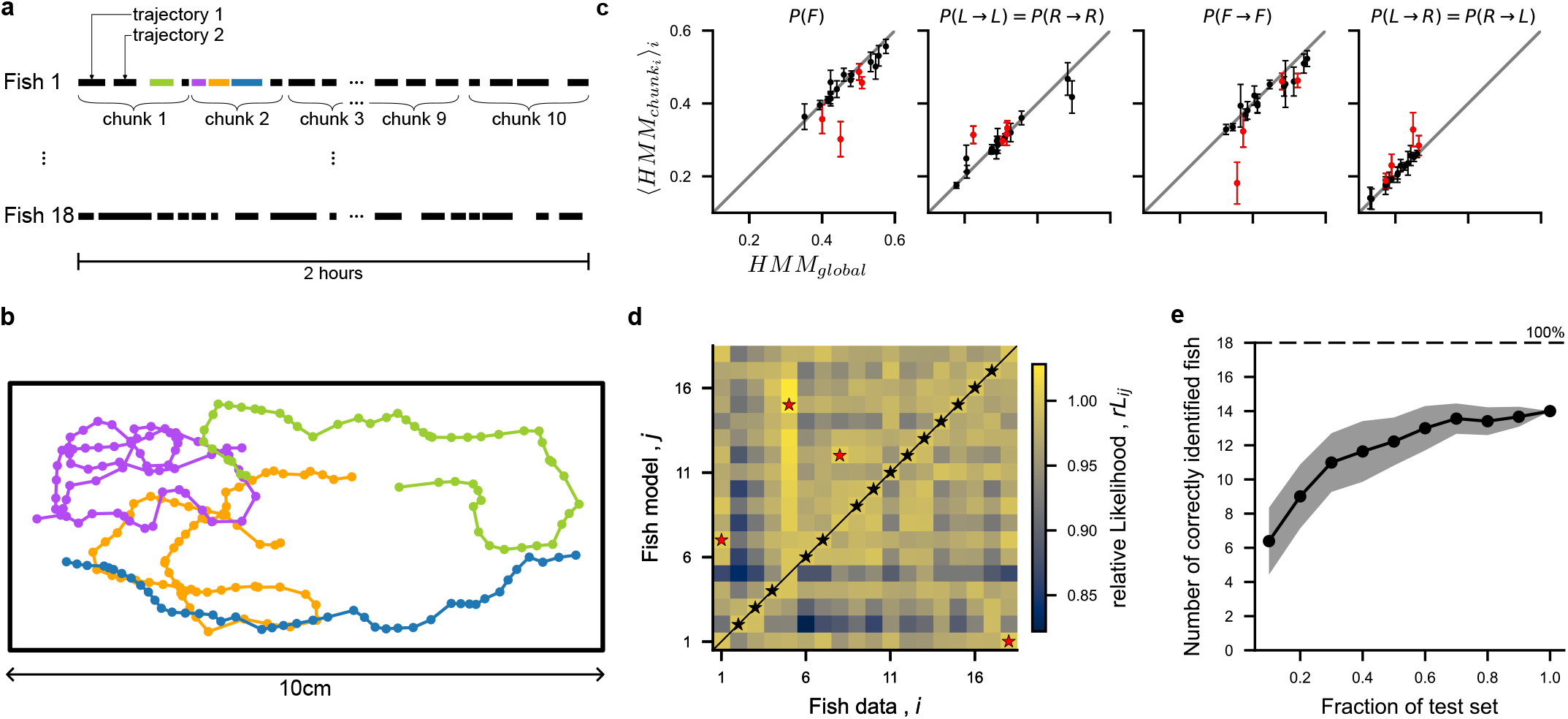
Fish identification from long trajectories: **(a)** Diagram describing the dataset. Trajectories from 18 fish recorded over 2-hour sessions, were each is split into 10 chunks (mean = 9.5 ± 0.5 trajectories per chunk) **(b)** Example trajectories for fish 1. **(c)** HMM parameters inferred from all the trajectories of a fish (referred to as global), compared with the HMM parameters trained on chunks of that fish’s trajectories. Only four HMM parameters are shown for clarity: the steady state probability of forward turns *P* (*F*), the transition probabilities for forward-forward *P* (*F → F*), turn-turn in the same direction *P* (*T*_1_ → *T*_2_ | *T*_1_ = *T*_2_), and turn-turn in opposite direction *P* (*T*_1_ → *T*_2_ |*T*_1_ ≠ *T*_2_) (see FigS5 for all parameters). Each dot represents a fish, and the error bars correspond to the standard error of the mean. Points labeled in red correspond to fish misidentified in panel d. **(d)** Confusion matrix between data coming from fish i and HMM trained on fish j. The relative likelihood 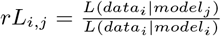 is used to evaluate which fish identity is most likely according to each model (indicated with black stars for correctly identified fish, and red stars for misidentification). **(e)** Number of correctly identified fish determined from model likelihood when only a fraction *f* of the test data is used for identification. The shaded area indicates the standard deviation across 100 trials. In each trial, the data trajectories of each fish were randomly split into train and test sets (50%).

These results suggest that the HMM models can be used to distinguish different fish from observations of their bout sequences. To test this hypothesis, we split the trajectories of each fish into a training and a withheld test set. After training the HMM on the train set for a particular fish, we computed the likelihood of all fish trajectories in the test set, and compared them. For 14 out of the 18 fish, the test set that yield the maximum likelihood rightly identifies the fish used to train the HMM (Fig.3d). This finding suggests that the HMM captures behavioral parameters which are distinctive enough to discriminate between different fish. Given the large variability exhibited by a single fish, one expects this discriminative ability to increase with the duration of the training sequences. To quantify this, we further evaluated the likelihoods of subsets of the test fish trajectories, and recorded the number of times that the maximum likelihood HMM corresponded to the correct fish (Figure 3e). Even when withholding 80% of the sequence, we were able to correctly identify 10 out of the 18 fish. These results suggest that individual fish exhibit variable but distinctive behavior which can be captured by the HMM.

### E. Modeling of neural data

The selection of turning bouts orientation in zebrafish is known to be controlled by a small bilaterally distributed circuit in the anterior hindbrain, called *Anterior Rhombencephalic Turning Region* (ARTR). This circuit displays self-sustained alternating activity between its left- and right-lateral sub-population, with a period of the order of tens of seconds (Fig.1e). The animal tends to execute left turns when the left ARTR is active while the right ARTR is inactive (and vice versa for right turns) [18].

In contrast, no specific circuit has yet been identified for the selection of turn vs forward bouts. The hypothesis that two distinct circuits are involved in bout-type selection is consistent with the 4 states Markovian model of navigation, in which two independent Markov chains drive the two selection processes. However, the 3-states Markovian model supported by the HMM analysis suggests that the same circuit (ARTR) could drive the selection of all 3 bout-types.

In order to test this hypothesis, we re-analyzed the ARTR recordings reported in Wolf *et al*. [27] using a 3-state HMM (Fig.4a). We posit an independent neural model for the activity of the N recorded neurons, yielding, for each state, the emission probability:

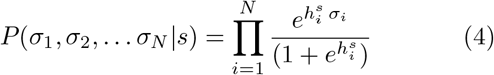

where (*σ*_1_, *σ*_2_, … *σ*_*N*_) is a neuronal configuration, s is the hidden state, and 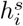 is the local field representing the effective excitability of neuron *i* in state *s*. The model thus includes 3 × N parameters 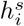, associated to each neuron and each hidden states. Notice that for the neural HMM, the non-handedness of the behavioral HMM is not enforced.

The distribution of fields 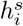 for the 3 hidden states, shown in Fig.4a, are used to assign labels to the three states (see Materials and Methods IV F). Consistent with our current understanding of the ARTR function for turn selection, the state with large values of the fields on the left and smaller values of the fields on the contralateral side is labeled “left” (and vice versa for “right”). The third state exhibits similar distributions of fields for neurons on the left and right side of the ARTR, and is labeled forward in analogy with behavior. The ARTR activity is thus modeled as a sequence of left-right-forward states.

With this classification, the forward state corresponds to a low mean neuronal activity of both the left (*m*_*L*_) and right (*m*_*R*_) sides of the ARTR, while turning states are associated with large activity on the ipsilateral side of the ARTR (left state : *m*_*L*_ > *m*_*R*_, right state : *m*_*L*_ < *m*_*R*_, see Fig.4b-d).

This model accurately captures the mean activity of each neuron (Fig.4f), as well as the pairwise correlations between contralateral neurons. However, ipsilateral pairwise correlations are not as well reproduced, showing lower covariance in the generated data (Fig.4f). This mismatch presumably comes from the fact that the activities of neurons within a state are uncorrelated in our emission probabilities, while recurrent interactions in the ARTR circuit produce correlations. These would be better modeled with emission probabilities including effective interactions between neurons [27].

### F. Comparison of Behavior and Neuronal HMMs

In the preceding sections, we demonstrated that both the reorientation behavior and the neuronal activity of the *Anterior Rhombencephalic Turning Region* (ARTR) can be effectively modeled using three-state Hidden Markov Models (HMMs). However, it remains unclear whether the three states identified in the Behavioral HMM (B-HMM) directly correspond to those inferred in the Neuronal HMM (N-HMM).

Unfortunately, there is currently no publicly available dataset offering simultaneous recordings of freely swimming larvae kinematics and neuronal activity, which would enable direct comparison of B-HMM and N-HMM states for individual bouts. Current research addressing this question largely relies on experimental paradigms where larvae are either paralyzed with electrophysiological recording of motor nerve signals (fictive swimming preparations)[18, 30, 31], or head-embedded with a free-moving tail (head tethered preparations)[32–35]. In fictively swimming preparations, whilst the classification of left-vs-right bouts is feasible based on the asymmetric nature of the motor command, such experiments lack the level of precision required to discriminate forward-vs-turning bouts [18]. On the other hand, head tethered preparations allow forward-left-right bout classification [32, 34], but typically rely on visual stimuli to elicit behavior [32–35] as the spontaneous sequence of bouts is strongly disrupted in comparison with freely swimming contexts [36].

We hereafter propose to circumvent these experimental challenges by comparing the statistical structures of the reorientation sequences inferred from the two datasets presented in sections II A and II E. The transition probabilities *P*(*s*_*n*_ → *s*_*n*+1_) obtained from B-HMM and NHMM at all recorded temperatures are shown in Fig.5b. Comparison of these transition rates require to first correct them for differences in sampling rates. Indeed, neural transition rates are computed from neuronal recordings performed at ∼ 6Hz (depending on the dataset, see Materials and Methods IV B), while for behavior, the sequences are divided into swim bouts triggered at an average rate of ∼ 1Hz, depending on the temperature.

To bridge the gap between neuronal and behavioral datasets, one needs to estimate how the behavior is subsampled from the neuronal activity. To do so, we computed the distribution of sojourn times Δ*t*_*s*_ of all three states in both B-HMM and N-HMM, where Δ*t*_*s*_ = *t*_*k*_ − t_1_ is the duration of a sequence (*s*_1_, …, *s*_*k*_) of k consecutive states s observed at times (*t*_1_, …, *t*_*k*_). We found the neuronal sojourn times to be significantly longer than the behavioral sojourn times (Fig.5a). The optimal temporal scaling factor *f*_*N/B*_ for which the distribution of neuronal sojourn times matches the distribution of behavioral sojourn times (see Materials and Methods IV G) was *f*_*N/B*_ ≈ 0.44. Interestingly, this value appears to be consistent with findings from Dunn *et al*. [18], which reported the mean interbout interval for fictive swimming to be 0.41 times slower than for freely swimming.

Using this temporal re-scaling factor, we find that the transition probabilities *P*(*s*_*n*_ → *s*_*n*+1_) for behavior and ARTR models are similar (RMSE = 0.1, see Fig.5b), indicating that the behavioral and neuronal state sequences share similar underlying structures. This is remarkable as the number and meaning of the neuronal internal states were not *a priori* fixed, but entirely assigned by N-HMM after training.

This result supports our hypothesis that the ARTR not only governs the selection between rightward and leftward turning bouts, but also controls the bout-type selection, forward *vs* turn. To test this claim further, we analyzed in more detail the statistics of trajectories in the bout space inferred from the ARTR dynamics and from behavioral data. We specifically examined the bout sequences leading to a change in orientation, such as transitions from *L* to *R* and vice-versa. Such orientational switches can be either direct, e.g. *L* → *R*, or may include an intermediate forward bout, *L* → *F* → *R* (Fig.5c). Using the ARTR signal, we found that the second path is strongly favored as evidenced by the the fact that 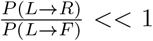. A comparable value of this ratio is observed in the behavioral data (Fig.5d), indicating that fish indeed tend to execute a forward bout when changing orientation. This statistical bias would be difficult to understand under the standard model that posit the existence of independent neural circuits governing orientation and bout-type selection, respectively. In contrast, in our model, it emerges naturally from the the phase space structure of the ARTR dynamics as shown in Figure 4c and Figure 5c. The L-Shaped distribution of {*m*_*l*_, *m*_*R*_} constrains the Left-to-right (or Right-to-Left) trajectories to pass through a symmetrical, low activity state, thus favoring intermediate forward bouts.

**FIG. 4.**
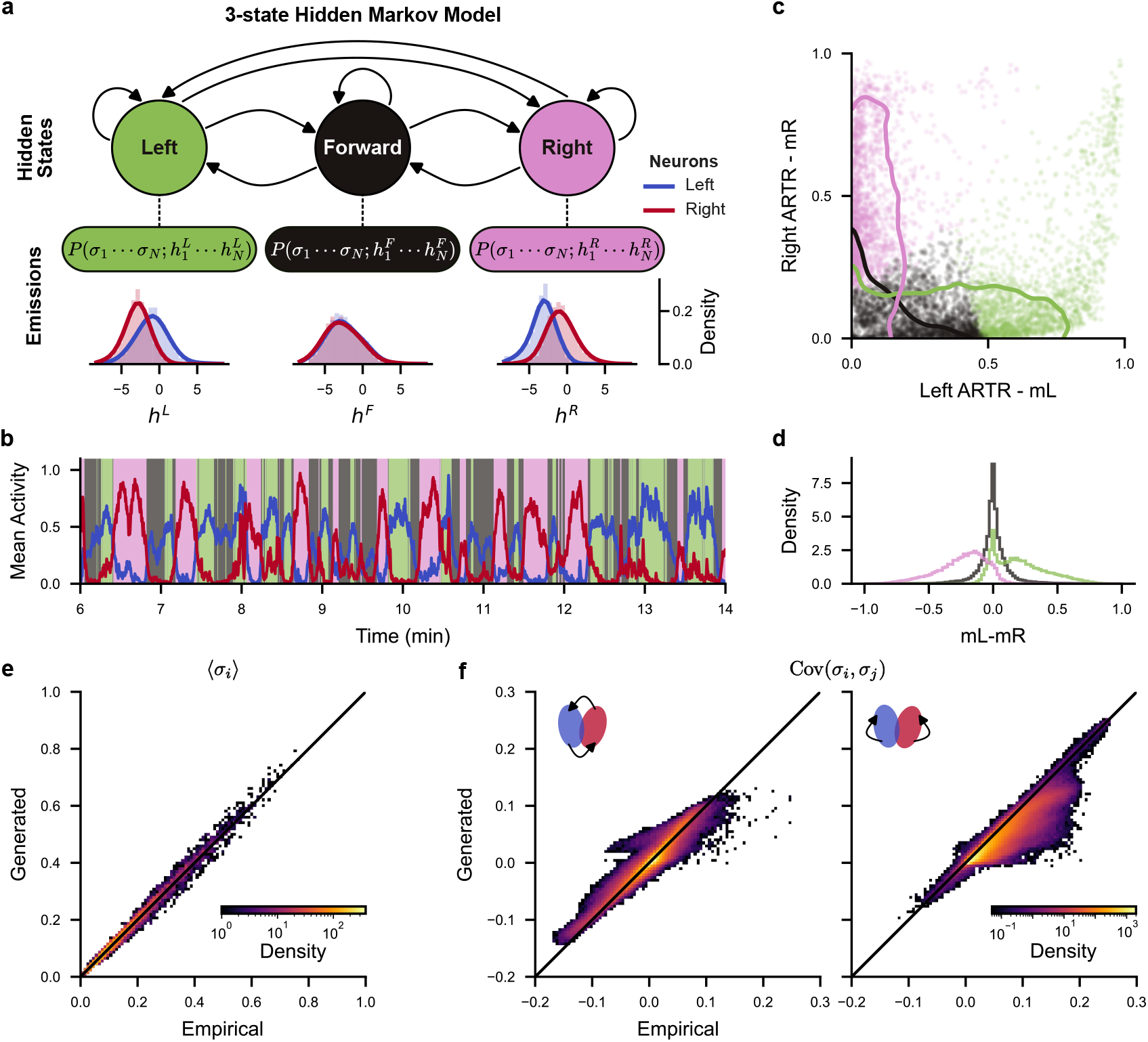
3-state Hidden Markov Model (HMM) describes ARTR neuronal statistics: **(a)** Diagram illustrating the 3-state Hidden Markov Model_*s*_(HMM) with emissions described as independent models of the ARTR neuronal population, see Eq. (4). Distributions of fields 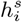 are shown for all recorded fish for neurons in the left and right ARTR (in blue and red respectively). **(b)** Example ARTR activity (see Fig. 1f) classified with the 3-state HMM. Blue and red lines represent the mean activity of neurons in the left (*m*_*L*_) and right (*m*_*R*_) ARTR, respectively. **(c)** HMM classification in the (*m*_*L*_, *m*_*R*_) space. Dots represent neuronal configurations taken from the example recording in panel b. Solid lines represent 90% of the distributions for all recordings combined. **(d)** Distributions of *m*_*L*_ − *m*_*R*_ for each hidden state and all recordings combined. **e-f** Comparison of empirical and HMM-generated neuronal statistics for all recordings combined. **(e)** Mean activity ⟨*σ*_*i*_ ⟩ of neuron **(f)** Covariance Cov(*σ*_*i*_, *σ*_*j*_) of neurons *i* and *j* on opposite sides (left plot) and on the same sides (right plot) of the ARTR.

**FIG. 5.**
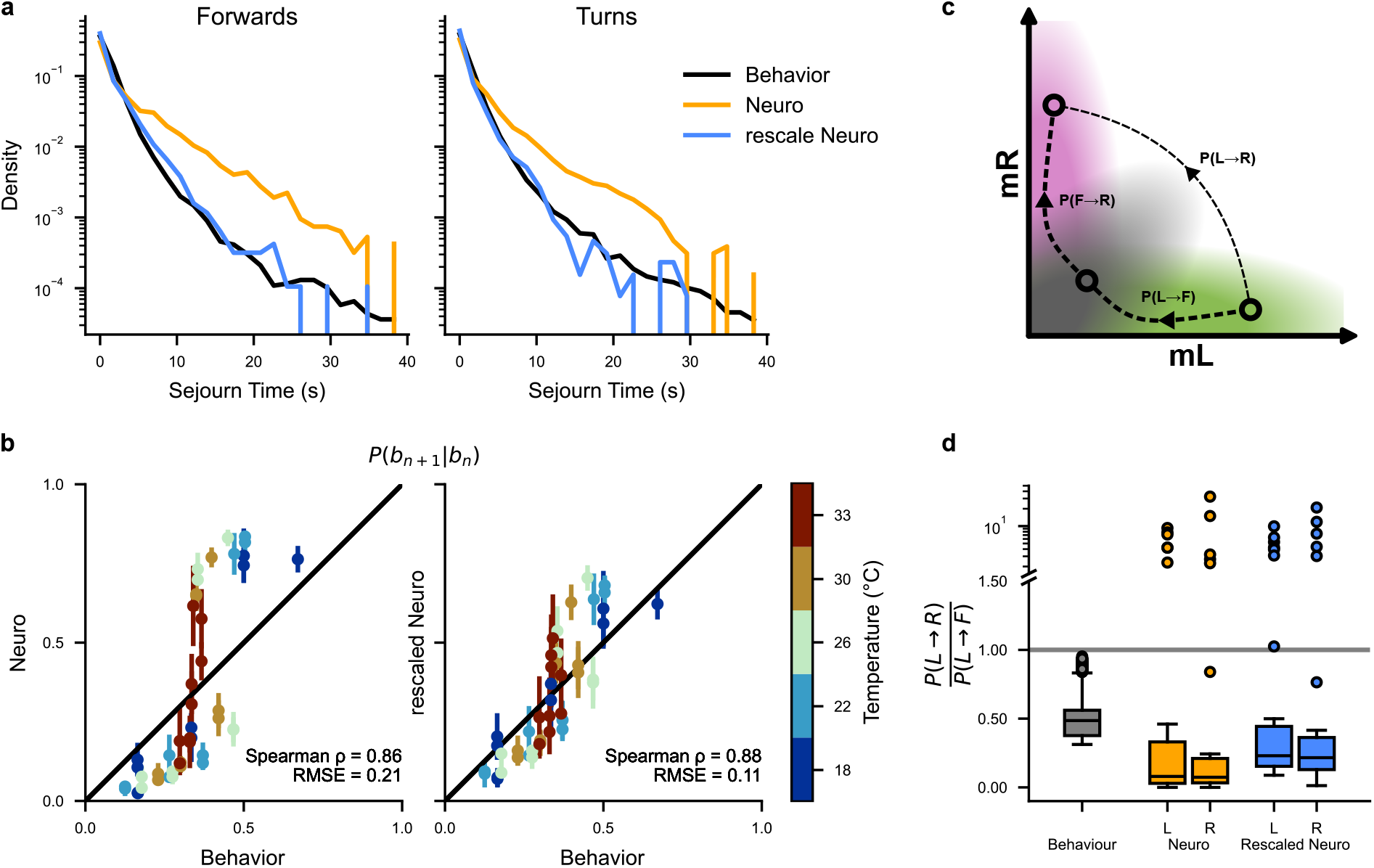
Behavior vs. Neuronal temporal structure: **(a)** Distribution of forward and turn sojourn times for the behavior (black) and neuronal data before (orange) and after temporal re-scaling (magenta). A single re-scaling factor is used for forward and turning states, for all temperatures, and for all recordings. **(b)** Comparison of behavior and neuronal state-transition probabilities *P* (*s*_*n*_ → *s*_*n*+1_), before (left plot) and after (right plot) temporal re-scaling. Each dot represents a single transition probability at a given temperature. For neuronal state-transition, the mean and standard error of the mean for all recordings at specific temperatures are shown. **(c)** Diagram showing two possible transition trajectories between left and right states in ARTR mean-activity space. Transitions through the forward state are more probable (see panel d). **(d)** Distributions of 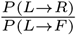 the behavior (black) and neuronal data before (orange) and after temporal re-scaling (magenta), with all temperatures combined. These distributions are depicted as standard box plots (median and quartiles), as well as outlier points lying further than 1.5× the inter-quartile range from the median.

### G. Generation of synthetic behavior with the neural model

Until now, we compared neuronal and behavioral data by examining only the short-scale statistical structures of the HMM-inferred state sequences. We now wish to test whether it is possible to compare full trajectories by leveraging the generative nature of the HMM. Specifically, we use the N-HMM model to generate long synthetic trajectories and compare their statistics with those of freely swimming fish. This approach allows us to assess whether the N-HMM, when combined with appropriate scaling and behavioral parameters, can reproduce the complex statistical properties of exploration at various temperatures.

#### 1. Generation of synthetic neural and reorientation trajectories

As stochastic processes, Hidden Markov Models (HMMs) can be sampled to generate new sequences of internal states. Following the previous section II F, we hypothesize that the internal states of a Neural HMM (NHMM) match the behavioral internal states, after proper temporal rescaling. Therefore, we expect that it should be possible to generate artificial swim trajectories from the N-HMMs.

Using the N-HMMs associated to individual fish recordings, we started by generating synthetic temporal sequences of neural states 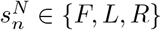. We then sampled the behavioral distribution of inter-bout intervals *δt*_*n*_, rescaled by the scaling factor *f*_*N/B*_ ≈ 0.44 obtained in the previous section II F. This simulates a stochastic bout-initiation process with the correct temporal characteristics, yielding synthetic sequences of bout internal states *b*_*n*_ for the behavior. For each state, we then sample the emission probability *E*(*δθ*_*n*_|*b*_*n*_) associated to the Behavioral HMM (B-HMM) inferred from all fish data to get a realization of the reorientation angle *δθ*_*n*_ (Fig. 6d). As expected, the distribution of these angles is in very good agreement with the ones observed in the behavioral data (Fig. 6a).

**FIG. 6.**
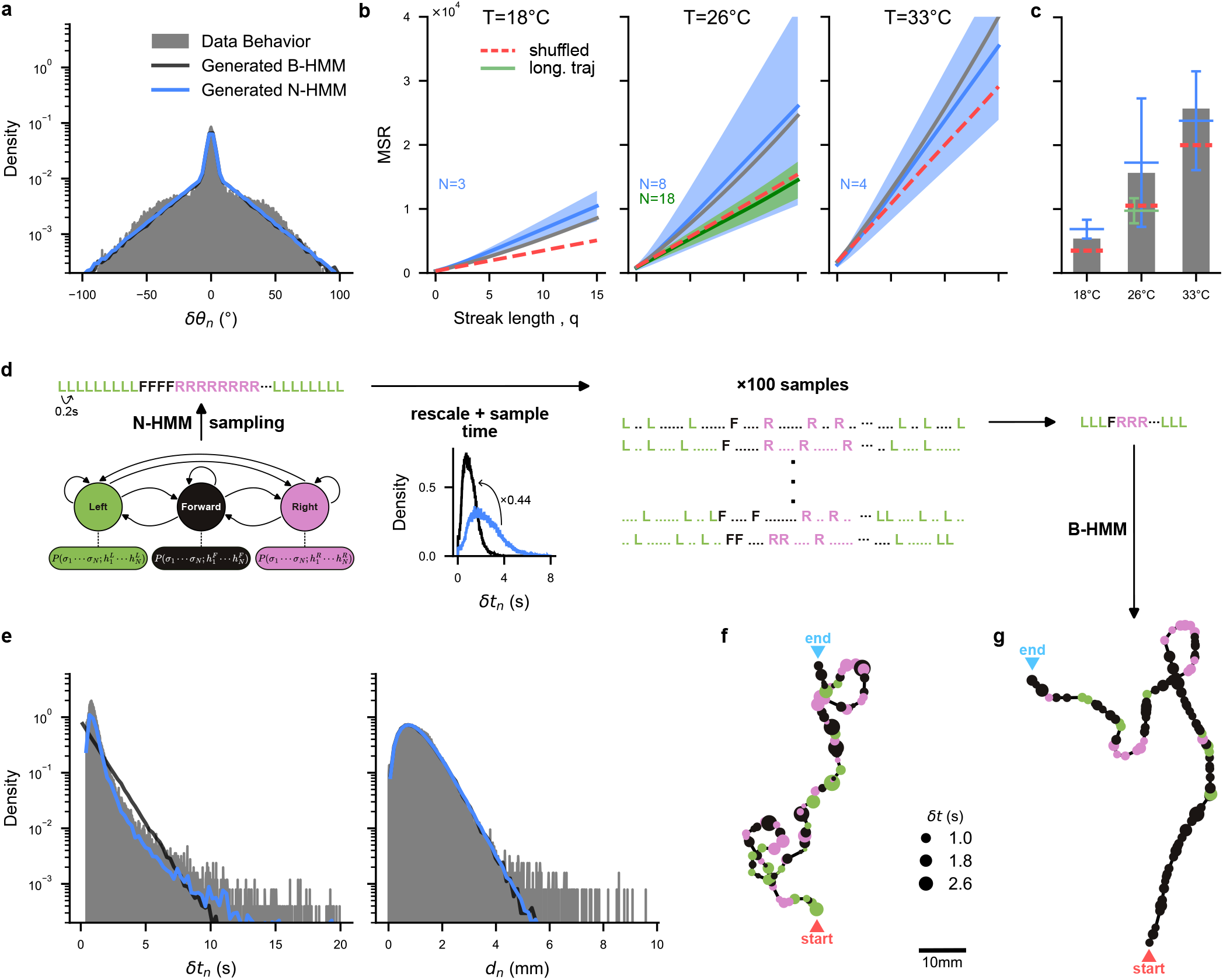
Generative ability of HMM models and trajectory reconstruction: **(a)** Distribution of bout angles *δθ*_*n*_ for the aggregated multiple-fish trajectories (gray), generated trajectories from Behavioral Hidden Markov Models (B-HMM; black), and generated trajectories from Neuronal Hidden Markov Models (N-HMM; blue), at 22°C. **(b)** Mean Square Reorientation (MSR) accumulated after *q* bouts for aggregated multiple-fish trajectories (grey), shuffled aggregated multiple-fish trajectories (red dashed), single-fish long trajectories (green), and trajectories generated from N-HMM (blue). For both long and N-HMM-generated trajectories, the mean and standard deviation over all individual fish are shown (respectively with a solid line and filled band). (see Fig.S7 for individual trajectories and all temperatures) **(c)** Bar plot for the MSR at *q* = 10 bouts, with mean (horizontal bars) and standard deviation (vertical bars). **(d)** Diagram explaining the conversion from N-HMM generated state sequences to swim trajectories. The N-HMM is first sampled to generate a sequence of forward, left, right internal states. Time is then re-scaled using the scaling factor identified in Fig 5, and bout sequences are sampled 100 times based on the interbout interval distribution. A swim trajectory is constructed for each bout sequence by sampling the bout distances *d*_*n*_ and inter-bout intervals *δt*_*n*_ emission distributions in the B-HMM. **(e)** Distribution of inter-bout intervals *δt*_*n*_ and bout distances *d*_*n*_ for the aggregated multiple-fish trajectories (gray), generated trajectories from B-HMM (black), and generated trajectories from N-HMM (blue), at 22^*o*^C. **(f)** Example trajectory generated from B-HMM at 26^*o*^C. Point color corresponds to bout type (left, right, forward), and point size corresponds to inter-bout interval. **(g)** Same as panel f for a N-HMM-generated trajectory at 26^*o*^C.

We further characterize the trajectories using the Mean Square Reorientation (MSR) after *q* bouts:

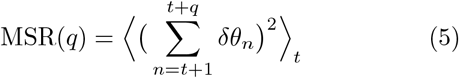

where the average is taken across all times and all trajectories.

Figure 6b shows the values of *M*_*q*_ obtained from NHMM-generated trajectories at different temperatures (see Fig.S7 for the remaining temperatures), as well as the MSR directly obtained from multiple-fish trajectories and long single-fish trajectories (only at 26°C).

We first notice that long individual fish trajectories at 26°C display large variability in MSR(q) values, compatible with the presence of fish-to-fish variability. This variability is washed out for the multiple-fish dataset (since individual trajectories are combined) providing a averaged MSR(q) for each temperature. Interestingly, the MSR for the long sequences of individual animals significantly differ from the MSR obtained from multiple-fish trajectories. This could be due to differences in experimental conditions, and in particular the effects of collective vs. isolated navigation [22].

N-HMM-generated trajectories have a MSR distribution compatible and encompassing the behavioral data in their variability. Such large variability is expected from the large fluctuations in neural brain states. Some trajectories generated with N-HMM show anomalously large angular persistence (see Fig.S7a), which may correspond to brain states where the *Anterior Rhombencephalic Turning Region* (ARTR) displays no left-right alternating behavior. This is expected, as the N-HMMs were established from spontaneous activity recordings of immobilized fish which were not constrained to swim-like behaviors. In Fig. 6c, we summarized and compared the results for the Mean Square Reorientation after 10 bouts, MSR(*q* = 10), for all temperatures. We found, consistently across temperatures, that the MSR of behavioral data are comprised within the one-standard-deviation confidence interval of N-HMM-generated trajectories.

As we show in the Appendix 1, the MSR of HMM-generated trajectories can be decomposed as the sum of a purely diffusive contribution, associated to the variance of bout angles, and of terms arising from time-correlations in bout type selection along a trajectory (see Eq. (A.28)). The increase of bout-angle variance with temperature is sufficient to explain the increasing trend of the mean MSR with temperature observed in Figure 6c (see Fig.S7).

#### 2. Generation of synthetic 2D trajectories

For the sake of completeness, we also used the N-HMM model to generate full synthetic 2D trajectories. To do so, for each bout state identified with the procedure reported above, we sampled an inter-bout interval duration *δt*_*n*_ and traveled distance *d*_*n*_ from their experimental distributions. We then reconstructed the coordinates of the virtual fish after *k* bouts, (*x*_*k*_, *y*_*k*_), through

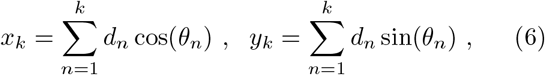

where 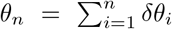 is the orientation angle of the fish at bout *i*, constructed as the cumulative sum of reorientation angles at previous bouts. An example trajectory is shown in Fig. 6g, and is qualitatively similar to the experimental counterpart recorded at the same temperature.

For comparison, we show synthetic trajectories generated from behavioral HMM. In practice, we expanded on the B-HMM introduced in section II B 2 by adding two new emission distributions corresponding to δt_*n*_ and d_*n*_. As done previously, the HMMs were first trained only on the re-orientation angles *δθ*_*n*_, before learning the emission distributions for *δt*_*n*_ and *d*_*n*_. We then plot the corresponding trajectories using Eq. (6), which are qualitatively similar to the N-HMM-generated ones, as illustrated by an example in Fig.6f. The similarity is quantified by the comparison of the distributions of bout angles, inter-bout duration intervals and traveled distances, see Fig.6f.

We report in Figure S7e-h, the outcome of an intermediate generative model, in which 2D swim trajectories are generated from experimental neural recordings. This is done by first identifying neural states from the recordings using the Viterbi algorithm, emiting inter-bout intervals using the same procedure as described above, and then feeding the resulting state sequences through the B-HMM.

## III. DISCUSSION

With the advancement of video-tracking and brain recording methods, behavioral neuroscience has changed radically in the last decade. It is now possible to study in great details animal behavior in unconstrained naturalistic conditions [37–39], while new recording methods give access to extended circuit activity encompassing several brain regions. Such experiments produce vast amounts of high-dimensional data, requiring automated yet robust and interpretable analysis methods.

An essential task is the identification of behavioral or neural states from the segmentation of the recorded time series, in order to extract low-dimensional representations that are easier to interpret. However, no definitive procedure exists for selecting the optimal number of states or for defining valid labeling criteria. This choice typically depends on available observables and involves a compromise between interpretability and accuracy of representation.

In our case, the difficulty stems from three main factors: (i) the dependence of swim bout kinematics with bath temperature, (ii) the inter-individual variability, and (iii) the overlapping distributions of reorientation angles for distinct bout types, in particular at low temperatures. Because they can accommodate such overlaps while taking into account the temporal regularities in the bout sequences, Hidden Markov Models (HMMs) are ideally suited for such a task. They are easily interpretable as the dynamics between the different internal states is described by a Markovian process. This makes HMM a powerful alternative to deep-learning-based methods, whose predictive power comes at the cost of interpretability.

In this study, we successfully applied a three-state HMM to parse behavioral and neural time series associated with exploratory dynamics. We showed that for behavioral data, HMM provided a less biased and more consistent method for bout-type labeling compared to standard threshold-based Markov Chain (MC) methods used in earlier studies.

This robustness proved essential as we investigated the effect of bath temperature on navigation. Zebrafish being cold-blooded animals, the water temperature is expected to directly affect muscle efficiency, leading to a systematic increase in the amplitude of reorientation elicited by bouts as temperature rises. When using hard-threshold-based MC methods, this may lead to a systematic but artifactual increase in the fraction of bouts labeled as turns with temperature. With HMM, this physiological effect of temperature is naturally accounted for through an adaptive adjustment of the effective threshold angle between turn and forward bouts. With this unbiased labeling, we found that the fractions of forward and turn bouts were only weakly dependent on temperature, in contrast with previously published analysis [22]. The primary effect of temperature of rising temperature is to progressively decrease bout-type persistence, i.e. the tendency of the animal to chain similar bouts. Interestingly, we found that all three bout types, and not just turns as previously reported, exhibit comparable persistence.

HMMs also demonstrated remarkable sensitivity to individual phenotypic variability. Inter- and intra-individual variability are ubiquitous traits of animal behavior [40, 41] and are necessary to ensure a trade-off between flexibility and adaptability to changing environmental demands and robustness in neural development [42, 43]. In Le Goc *et al*. [22], inter-individual differences were demonstrated on the same dataset using multiple kinematic parameters (including inter-bout interval, forward travel distance or reorientation amplitude). In contrast, our study shows that HMM can identify individual fish solely based on the dynamics of bout-type sequences. Moreover, HMM provides explicit likelihood evaluation for bout sequences for various individual-specific models, providing a quantitative measure of phenotypic proximity between animals or across time.

Since our approach is based on gait phenotyping and is independent of image features, it is compatible with low-resolution videos (in which only the animal’s position and orientation can be accessed) while still keeping versatility, reliability, and fast execution. This opens new opportunities for studying phenotypic variation in swimming behavior, potentially uncovering subtle effects on behavior of genetic, developmental, or environmental cues. This ability to precisely capture behavioral variability might also prove fruitful in order to explore the neural basis of individuality.

The fact that the fish directional dynamics can be-described by a three-state Markovian sequence, suggests that bout-type selection is likely governed by a single circuit, with the ARTR being the most plausible candidate. Since its discovery in 2012 [30], the ARTR has been viewed as a direction-selection hub, controlling lateralized behaviors such as tail flick and ocular saccade orientation [18, 44]. It also responds to lateralized visual stimuli, including binocular contrast and whole-field lateral motion [20, 44].

In the present study, we showed that a three states HMM can accurately describe ARTR neuronal data, and that this model is structurally and temporally similar to behaviorally-trained HMMs. This result suggests that the ARTR may also govern forward bout selection, unifying the control of all directional bout types within a single circuit. This interpretation is reinforced by the generation of synthetic, neuronally driven swimming sequences that closely matched the statistics of observed trajectories.

Bout-type persistence, as observed in behavioral assays, is mirrored in the slow sequential exploration of the three hidden states identified in neural recordings of the ARTR. Although the HMM enables the identification of these neuronal states, they provide no interpretation of how they emerge from interactions among the ARTR neuronal population. In fact, our implementation of HMM assumes the activity of neurons to be independent of each other when conditioned to a state.

In a recent study, we trained data-driven graphical models (Ising model) on ARTR activity sequences [27]. The Ising model uses activity patterns to learn the interactions between neurons but, unlike HMM, it ignores any temporal information in the data. Interestingly, the inferred Ising models tended to display three metastable states, two with high activity on either side and one “equilibrated” state with intermediate, balanced activity on both sides, consistent with the three hidden states found with HMM. This convergence underlines the complementary strengths of state-space and energy-based models in elucidating neural dynamics. While the former might enable capturing the temporal structure in collective neural activity, the latter offer insights into the underlying network interactions driving these states, and how metastability emerges within neural populations.

The exact mechanism through which the ARTR controls bout selection remains unclear. However, our findings suggest that ARTR subpopulations (right and left) might inhibit contraversive bouts (*i*.*e*. when the left side is active, it suppresses rightward turns, favoring leftward swim). In the equilibrated state, both inhibitory signals suppress turn bouts, leaving forward movement as the only option. Such a motor suppression mechanism is consistent with observations by Dunn *et al*. [18], which showed a continuous relationship between the lateral difference in the ARTR activity and the mean reorientation angle of the executed bout. While our model strongly supports this hypothesis, definitive experimental validation is required.

Last of all, to facilitate the accessibility and adoption of Hidden Markov Model (HMM) formalism for analyzing behavioral sequences, we provide a comprehensive and instructive Python tutorial (https://github.com/EmeEmu/IBIO-Banyuls2023-Python). This tutorial can be adapted for specific datasets or used as a resource for broader educational goals.

## IV. MATERIALS AND METHODS

### A. Behavioral Datasets

The behavioral dataset used in the present study is derived from Le Goc *et al*. [22], and can be accessed directly at https://doi.org/10.5061/dryad.3r2280ggw. This dataset comprises spontaneous swimming trajectories of 5 to 7 dpf zebrafish larvae, collected at controlled bath temperatures of 18°C, 22°C, 26°C, 30°C, and 33°C. A camera was used to continuously record the swimming behavior of the fish within an arena of 100×45×4.5mm^3^ for 30 minutes at 25 frames/second. To eliminate border effects, a Region of Interest (ROI) was defined at a distance of 5mm from the arena’s walls. Fish that swam outside the defined tracking ROI were considered lost, and a new trajectory was initiated upon their re-entry into the ROI. The identity of the fish is thus lost each time it exits the ROI. Therefore, the dataset contains a varying number of fish trajectories, ranging from 532 to 1513 trajectories across the different temperatures (mean = 1148). Individual trajectories were tracked offline using the open-source FastTrack software [45], and were then discretized into sequences of swimming bouts.

Each trajectory consists of a sequence of swim bouts, spanning from 9 to 748 bouts per trajectory (mean=60, distributions shown in Fig.S1a). From this extensive dataset, we primarily utilized the re-orientation angles, defined as the difference between the heading direction at bout *n* + 1 and the heading direction at bout *n*:

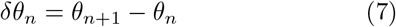

(a graphical illustration of this definition can be found in Fig.1c). This parameter encapsulates the angular change between consecutive bouts, providing insight into the fish’s ability to modify its orientation during swimming. We also used the interbout interval *δt*_*n*_ = *t*_*n*+1_ − *t*_*n*_ representing the elapsed time between 2 consecutive bouts, and the traveled distance 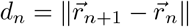.

On top of these multi-fish trajectories, we used in sections II D and II G a second dataset from Le Goc *et al*. [22] consisting in single-fish recordings. For this dataset, each fish (N=18) is placed alone in the arena at 26°C, and is recorded for 2 hours. With this experimental paradigm, the identity of the fish is conserved across trajectories, even when the fish leaves and re-enters the ROI.

### B. Neuronal Datasets

The neuronal dataset used in the present study is derived from Wolf *et al*. [27], and can be accessed directly at https://gin.g-node.org/Debregeas/ZF_ARTR_thermo. This dataset contains 32 one-photon Light-Sheet Microscopy recordings of spontaneous brain activity, for 13 zebrafish larvae (5 to 7 dpf) at 18°C, 22°C, 26°C, 30°C, and 33°C. It focuses on neurons from the *Anterior Rhombencephalic Turning Region* (ARTR), with ∼ 300 neurons (mean 307, std 119), recorded during ∼20min (mean 23, std 4 min) at ∼ 6Hz (mean 5.9, std 2.1 Hz).

### C. Emission of reorientation angles in the Hidden Markov Model

To validate the hypothesis that the re-orientation angles can be modeled using normal and gamma distributions, we compared the distribution of the data with a Gaussian Mixture Model (GMM) and a Gaussian&Gamma Mixture Model:

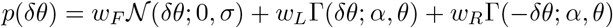

where *w*_*F*_ +*w*_*L*_+*w*_*R*_ = 1, and *w*_*F*_, *w*_*L*_, and *w*_*R*_ denote the weights for forward, left, and right states, respectively.

Using Quantile-Quantile (QQ) plots, we show that this last mixture model accurately reproduces the observed distribution of *δθ*_*n*_ in the data, and is much better than a GMM, especially in the tails of the distributions (Fig. S2a). We also confirmed that, once trained, the emission distributions do indeed match the observed reorientation distributions (Fig. S2b-c).

### D. Stubbornness factor

The *stubbornness* factor *f*_*q*_ is a measurement of the animal’s preference towards turning in the same direction over changing direction, after *q* intermediary forward bouts (Fig.S4c), as defined in (3).

It can be computed from a sequence of classified bouts *b*_*n*_ by first identifying and counting the q-plets *T*_1_ → *F*^*q*^ → *T*_2_ where *T*_1_ = *T*_2_ and where *T*_1_ ≠ *T*_2_:

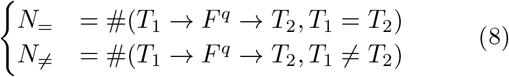

and then computing their ratio:

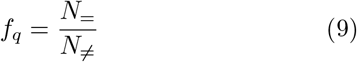

In practice, this ratio has a physical interpretation only for long sequences of bouts where *N*_=_ >> 1 and *N*_≠_ >> 1. As the trajectories in our dataset can be quite short (Fig. S1a), we compute *f*_*q*_ from all trajectories at a specific temperature, increasing the chance of observing a high number of stubborn (*N*_=_) and non-stubborn (*N*_≠_) trajectories.

By considering that the probability of a given q-plet is stubborn follows a binomial distribution (𝔼 (*N*_=_) = *pN* and 𝔼 (*N*_≠_) = (1 − *p*)*N* with *N* = *N*_=_ + *N*_≠_), we can evaluate the uncertainty in *stubbornness* as:

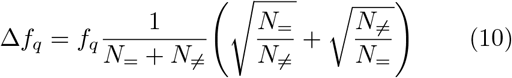

It is to be noted that these uncertainties are conservative estimates, as there exits a bias inherent to the dataset. Indeed, a very stubborn fish will tend to stay longer within the Region Of Interest (ROI) of the camera, leading to longer trajectories and therefore weighing more on the final result. Hence, it is unclear whether a *stubbornness* factor *f*_*q*_ = 1 ± 0.2 is truly significant (as suggested by the estimated error bars on Fig.S4d).

Furthermore, as the stubbornness factor is computed from all trajectories (and thus all fish) at a particular temperature, it represents an average behavior rather than an individual fish.

### E. Stubbornness factor and 3-state Markov Chain

The *stubbornness* factor can be defined directly from the transition matrix.

*For q* = 0, calculations are simple:

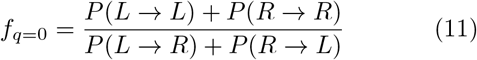

*For q* ≥ 1, the *stubbornness* factor is defined from the transition matrix as:

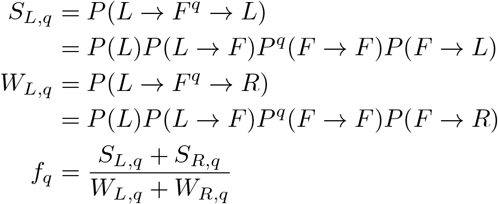

with *S*_*L,q*_ the probability of a trajectory which starts and ends with a left bout, *W*_*L,q*_ the probability of a trajectory which starts with a left bout and ends with a right bout, and *S*_*R,q*_ *W*_*R,q*_ their symmetrical opposites.

For a 3-state model, the forward-forward bout probability cancels out, giving:

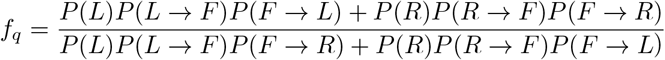

and with our non-handedness hypothesis: *P*(*L*) = *P*(*R*), *P*(*L* → *F*) = *P*(*R* → *F*), and *P*(*F* → *L*) = *P*(*F* → *R*), yielding:

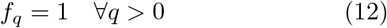

### F. Labeling of states in the neuronal Hidden Markov Model

The internal states of the Hidden Markov Models (HMMs) trained from neuronal activity are not *a priory* assigned to the Left, Right and Forward labels, and must therefore be re-ordered post-training.

We expect a certain symmetry in the system, where neurons in the left side of the ARTR will be more active during a Left state (and vice versa). Hence, we can use the excitability 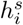 of neuron *i* in each internal state *s*, as defined in the emission distribution of the HMM (see Eq. 4). We define the lateralized excitability:

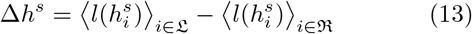

Where 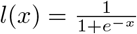 is the standard logistic function, and L and R are the sets of neurons located respectively in the left and right side of the ARTR. We thus label the HMM states such that

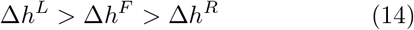

with *F, L*, and *R* the Forward, Left and Right internal states.

### G. Temporal re-scaling

To find the temporal re-scaling factor *f*_*N/B*_ between behavioral and neuronal models, we first compute the distributions of sojourn times Δ*t*_*s*_ for all states *s* ∈ {*F, L, R*} in both behavioral and neuronal Hidden Markov Models.

We then find the optimal re-scaling factor *f*_*N/B*_ for which the combined distributions 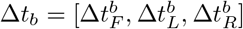 and 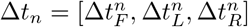 are as close to each other as possible :

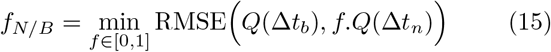

where *Q*(*D*) is the quantiles of a distribution *D*, and RMSE(**x, y**) is the Root Mean Squared Error between vectors **x** and **y** (see Fig.S6a).

For Markov chains, the transition matrix *P* = *P*(*s*_*n*_ = *s* → *s*_*n*+1_ = *s*′) represents the probability of transitioning in one step from state s to state *s*′. The transition probability *s* → *s*′ in k ∈ ℕ_1_ steps *P*(*s*_*n*_ = *s* → *s*_*n*+*k*_ = *s*′) is then the matrix power *P*^*k*^.

In order to apply the temporal re-scaling *f*_*N/B*_ between behavioral and neuronal models, we can thus compute the re-scaled transition matrix:

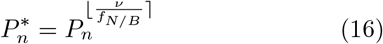

where *P*_*n*_ is the transition matrix inferred from neuronal data recorded at a frequency ν Hz.

### H. Mean Square Reorientation

To characterize the orientational diffusivity of the trajectories, we use the Mean Square Reorientation (MSR) accumulated after q bouts, as defined in equation (5) [20]. For infinitely large datasets with no left-right bias, we expect a centered distribution of reorientation angles ⟨*δθ*_*n*_⟩ _*n*_ = 0. However, this is not the case, particularly for the neuronal dataset where experimental limitations can induce strong biases. In particular, two of those limitations are due to the one-sided illumination in our Light Sheet Fluorescence Microscope [46]. First, due to scattering, the illumination beam is not uniform left-right across the brain, which can induce biases in the detection of neurons and their activity. Second, the non-uniform perception of light by the zebrafish larvae can elicit a phototaxis response, which is known to bias the activity of the *Anterior Rhombencephalic Turning Region* (ARTR) [44].

Since a non-zero bias can result in a distortion of the MSR (see Appendix 1), the MSR is computed from 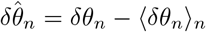 instead of *δθ*_*n*_.

## Acknowledgment

We acknowledge the following funding:

| Author | Funder |
| --- | --- |
| M. DK. | École Doctorale Frontière de l'Innovation en Recherche et Education - Programme Bettencourt |
| J. FdCD. | Université PSL, AI Junior Fellow program |
| M. C. | European Union, Horizon 2020 Programme (H2020 MSCA ITN Project SmartNets GA-860949) |
| V. B. | European Research Council (ERC) under the European Union's Horizon 2020 research innovation program grant agreement number 715980 |
| R. M., G. D., S. C. | Locomat ANR-21-CE16-0037 |

## Data and Code availability

All the data and code used in the present article are available under GNU General Public License version 3 at https://github.com/ZebrafishHMM2023/ZebrafishHMM2023_CodeAndData/tree/bioRxiv.

The custom Julia implementation of Hidden Markov Model used is available under MIT License at https://github.com/ZebrafishHMM2023/ZebrafishHMM2023.jl/tree/bioRxiv.

We also provide two tutorials for the use of Hidden Markov Models for behavioral sequence analysis. The first one was created for the Cogmaster “Machine learning for cognitive science” course, and is available at https://github.com/CoccoMonassonLab/ZebrafishHMM. The second one was created for the i-Bio Summer School ”Advanced Computational Analysis for Behavioral and Neurophysiological Recordings” held in Banyuls-sur-Mer in the summer of 2023, and is available under GNU General Public License version 3 at https://github.com/EmeEmu/IBIO-Banyuls2023-Python.

## APPENDICES

### 1. Mean squared reorientation

The mean-square reorientation (MSR) of a trajectory at lag *q* is defined as [20]:

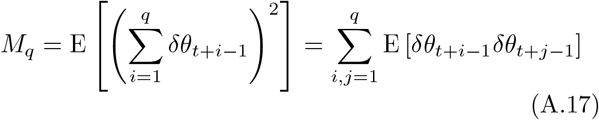

where it is assumed that E[*δθ*] = 0. The average is taken over time *t*. Assuming equilibrium, this is independent of *t*, and should depend only on the separation |*i* − *j*|,

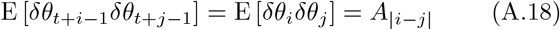

where A_|i−j|_ stands for the time equilibrated autocorrelation function:

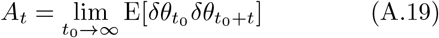

In particular *A*_0_ = E[*δθ*^2^] is just the variance of *δθ*. It follows that,

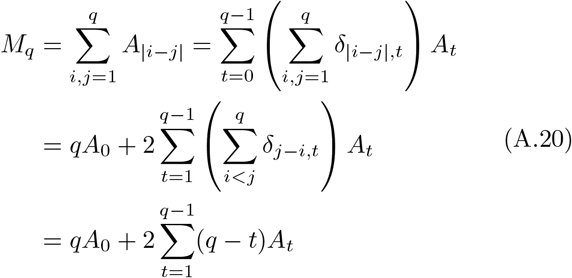

Note that for a random walk without any correlations across time, *A*_*t*_ = 0 for t > 0. In this case, *M*_*q*_ = *qA*_0_ grows linearly with *q*.

On the other hand, it is expected that A_*t*_ → 0 as *t* → ∞, and usually this decay is exponentially fast in time. Therefore, for large q, we get the following asymptotic expression for *M*_*q*_:

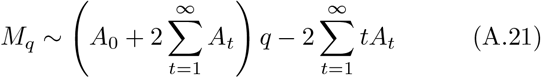

Notice that this is affine in q, with the coefficient 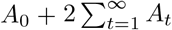. Therefore, *M*_*q*_ is initially linear in q with slope *A*_0_ for small *q*, then has an elbow and eventually approaches the asymptotic slope 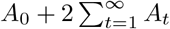 as *q* → ∞. This asymptotic slope is different from A_0_ only if the process exhibits non-trivial autocorrelations in time.

#### a. MSR for the HMM

As an illustration, we can compute all these quantities exactly for the HMM as follows. For the autocorrelation, we have:

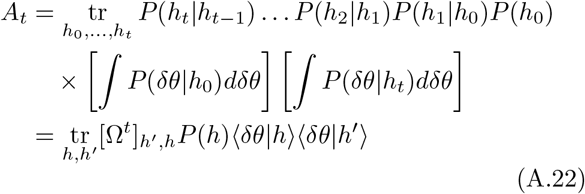

where [Ω]_*h*_*′,h* = *P*(*h*′ | *h*) is the transition matrix of the HMM. We will assume here that the initial state is sampled from *P*(*h*) = *p*_eq_(*h*), the equilibrium distribution of hidden states of the HMM, which satisfies the stationarity equation:

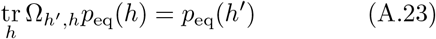

Note also that E[δθ] = 0 implies that _*h*_ p_eq_(h)⟨δθ|h⟩ = 0. Now let p_1_(h), …, p_*L*_(h) denote the remaining eigenvectors of Ω, with the associated eigenvalues λ_1_, …, λ_*L*_. By the Perron-Frobenious theorem, these remaining eigenvalues are all smaller than one in absolute value. The vector P(h) ⟨δθ | h⟩ can be writen in the basis of this eigenvectors,

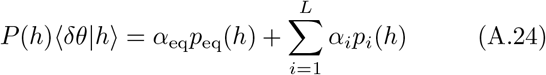

for some coefficients *α*_eq_, *α*_1_, …, *α*_*L*_. Then it follows that,

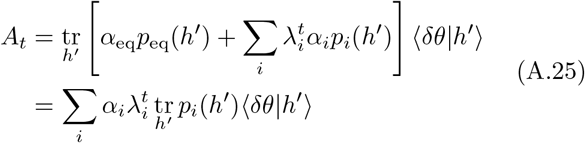

Since the |λ_*i*_| < 1 it follows that *A*_*t*_ → 0 exponentially fast as *t* → ∞. Moreover we can compute,

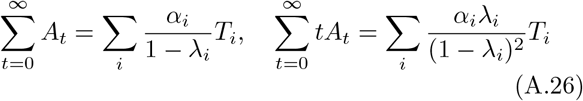

where

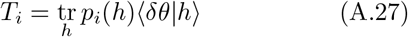

These expressions then give a complete and exact characterization of the MSR for the HMM.

#### b. Standardized MSR

**FIG. S1.**
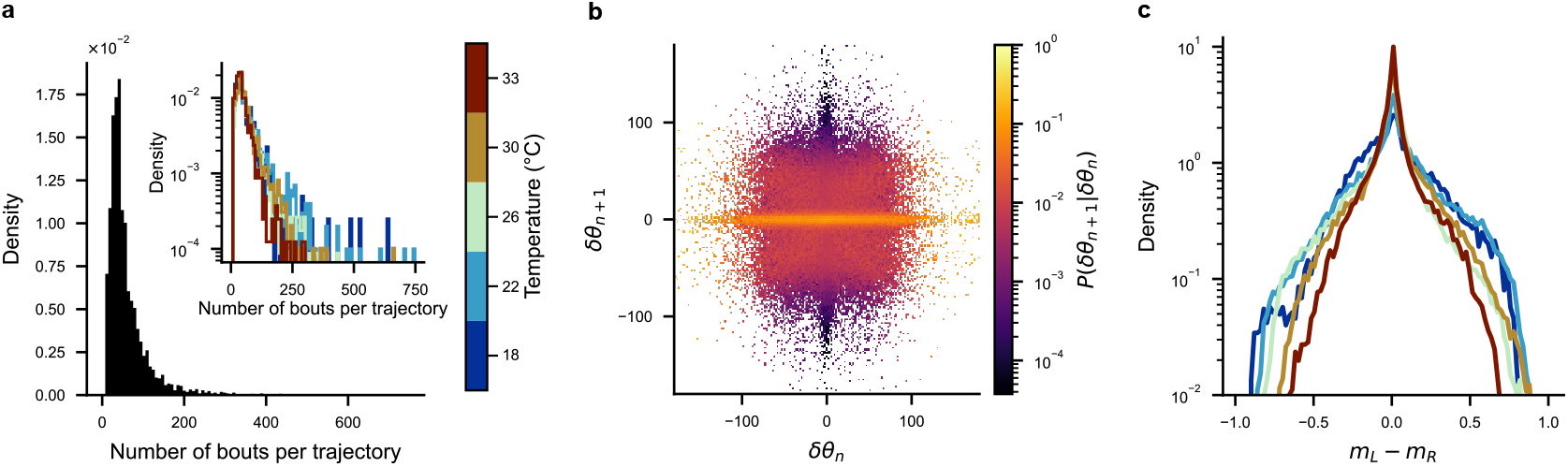
Supplementary panels to Fig.1: **(a)** Distributions of the number of bouts per trajectory in the entire behavioral dataset (black), and for each recorded temperature (inset, colored). **(b)** Observed transition probabilities between reorientation angles for the entire behavioral dataset. **(c)** Distributions of the difference between mean activities in the left (*m*_*L*_) and right (*m*_*R*_) *Anterior Rhombencephalic Turning Region* for all fish at each recorded temperature.

The MSR as defined in Eq. A.17 includes both the diffusive contribution from the initial term A_0_ and contributions arising from non-trivial time correlations in the process coming from the terms *A*_*t*_ for *t* > 0. As already pointed out, this initial term *A*_0_ = E[*δθ*^2^] is just the variance of the distribution of bout angles and is insensitive to time correlations. To emphasize the time correlations we may normalize the trajectories by defining:

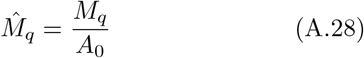

By comparing with Eq. A.21, we see that 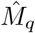 has initially a slope ≈ 1 for small *q*, then has an elbow_L_and eventually approaches the asymptotic slope 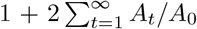 for large *q*.

In contrast to 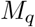, the quantity 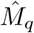 is better suited to compare the time correlations of very diverse trajectories because it is insensitive to variations of E[*δθ*^2^]. Figure S7c-d plots the normalized MSR from Eq. (A.28) for the various trajectories and temperatures considered before in Figure S7c-d. We observe that the standardized MSR exhibits comparable behavior across various temperatures, suggesting that the trend of the unnormalized MSR observed in Figures 6b-c and S7a-b is just due to an increase in the bout angle amplitudes E[*δθ*^2^] with temperature, but not due to changes in the structure of their time correlations.

**FIG. S2.**
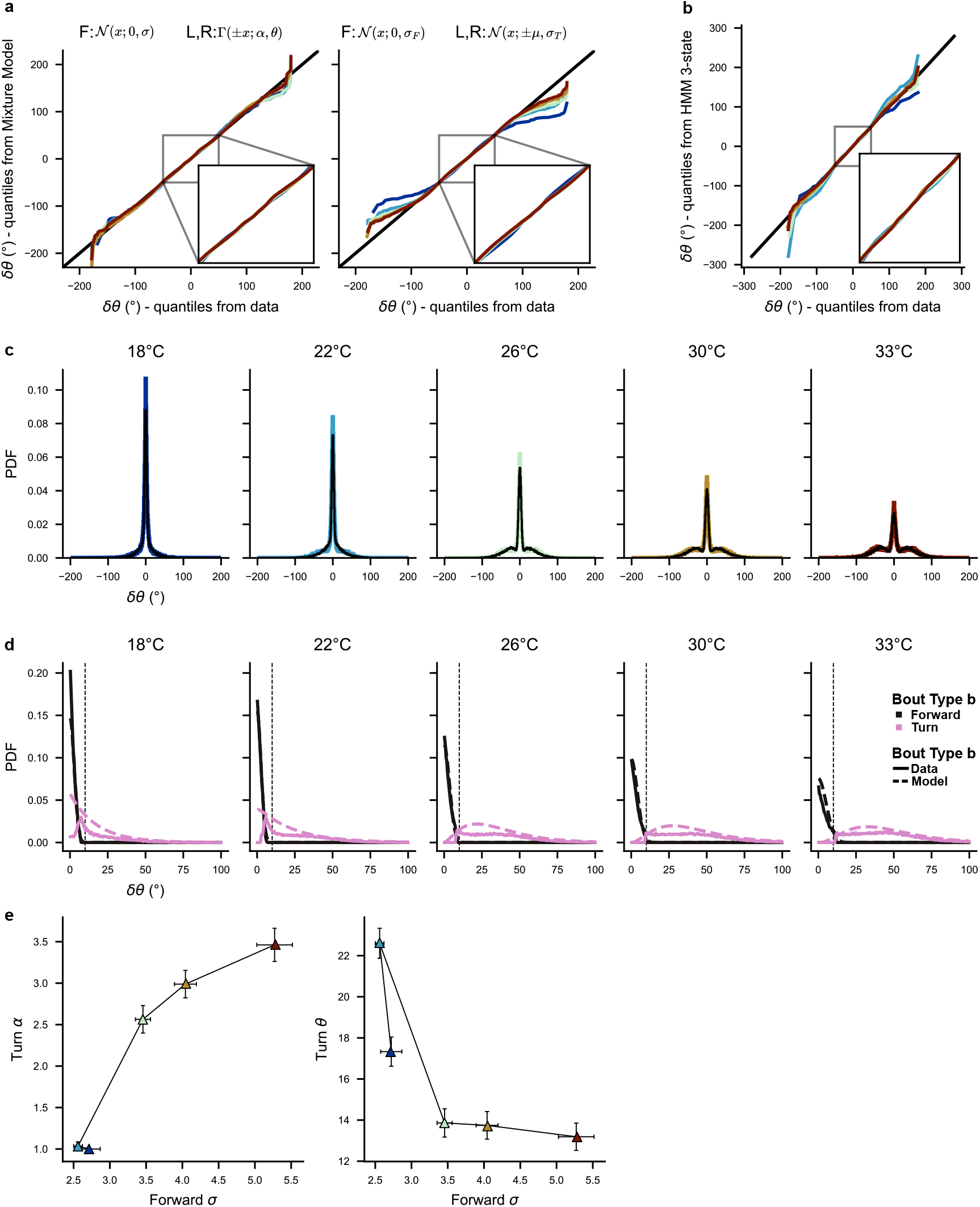
Supplementary panels to Fig.2 - Emission distributions: **(a)** Quantile-Quantile plot between distributions of reorientation angles observed from the data and Mixture Models, at each temperature. *Left:* Mixture Model defined from a central Normal distribution (forward bouts) and two Gamma distributions (left and right turning bouts), corresponding to the model of HMM emissions. *Right:* Gaussian Mixture Model. *Insets:* Zoom on ±50°. **(b)** Quantile-Quantile plot between the distributions of reorientation angles observed from the data and the distributions of reorientation angles generated by HMM. *Insets:* Zoom on ±50°. **(c)** Comparison between the distributions of reorientation angles observed from the data (colored) and the distributions of reorientation angles generated by the 3-state Hidden Markov Model (HMM; black), for each temperature. **(d)** Distributions of absolute reorientation angles labeled as forward bouts (solid black) and turning bouts (left or right; solid pink) by the Hidden Markov Model (HMM). Dashed lines show the HMM emission distribution for forward and turning bouts (black and pink respectively). The threshold *δθ*_0_ = 10° used in the Markov Chain model is shown for reference as a vertical black line. **(e)** Parameters of the HMM emission distribution, with *σ* the standard deviation of the central Normal distribution (forward bouts), *α* and *θ* the shape and scale of the Gamma distribution (turning bouts). Each dot corresponds to one temperature, and error bars were computed from the minimum-maximum of 100 cross-validations (trained on randomly selected 50% of the datasets).

**FIG. S3.**
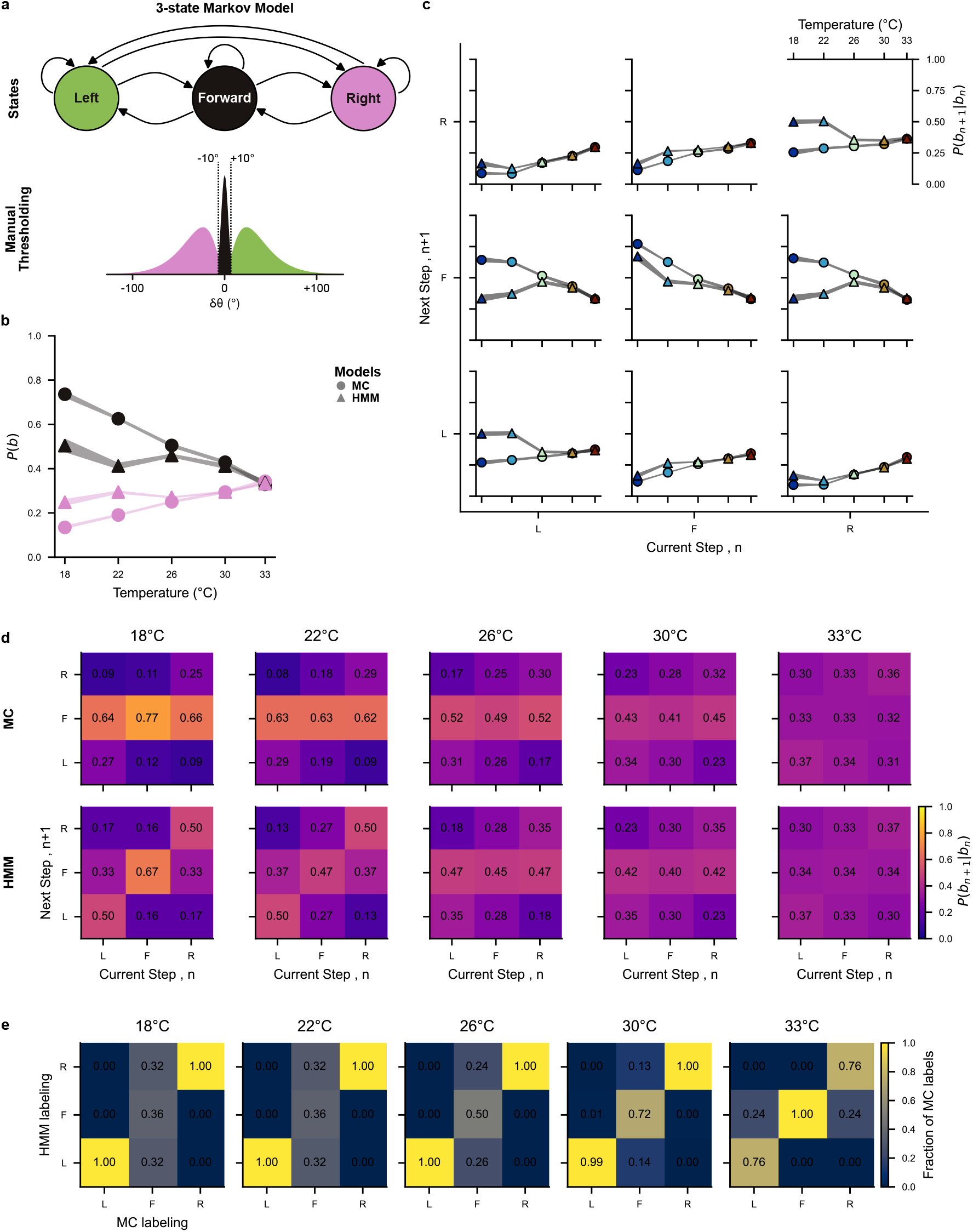
Supplementary panels to Fig.2 - Comparison between Markov Chain and Hidden Markov Model: **(a)** Diagram illustrating the 3-state Hidden Markov Model (HMM) with emissions modeled as a normal distribution for forward bouts and gamma distributions for turning bouts. **(b)** Temperature dependence of the steady state bout probabilities *P* (*s*) for forward bouts (*s* = *F*, black) and turning bouts (*s* ∈ *L, R*, pink), for both the Markov Chain inferred from thresholded reorientations (MC, circles) and Hidden Markov Models (HMM, triangles). **(c)** Temperature dependence of the transition probabilities *P* (*s*_*n*_ → *s*_*n*+1_) between forward (F), left (L), and right (R) bouts, for both the Markov Chain (MC, circles) and the Hidden Markov Model (HMM, triangles). **(b**,**c)** The width of the shaded curves represent the minimum-maximum of 100 cross-validations of both models inferred from randomly selecting 50% of the data. **(d)** Transition matrices between forward (F), left (L) and right (R) states, for both the Markov chains inferred from thresholded data (MC) and Hidden Markov Model (HMM), and for each temperature. **(e)** Confusion matrices between labeling of MC and HMM for all temperatures (normalized with respect to the MC labeling).

**FIG. S4.**
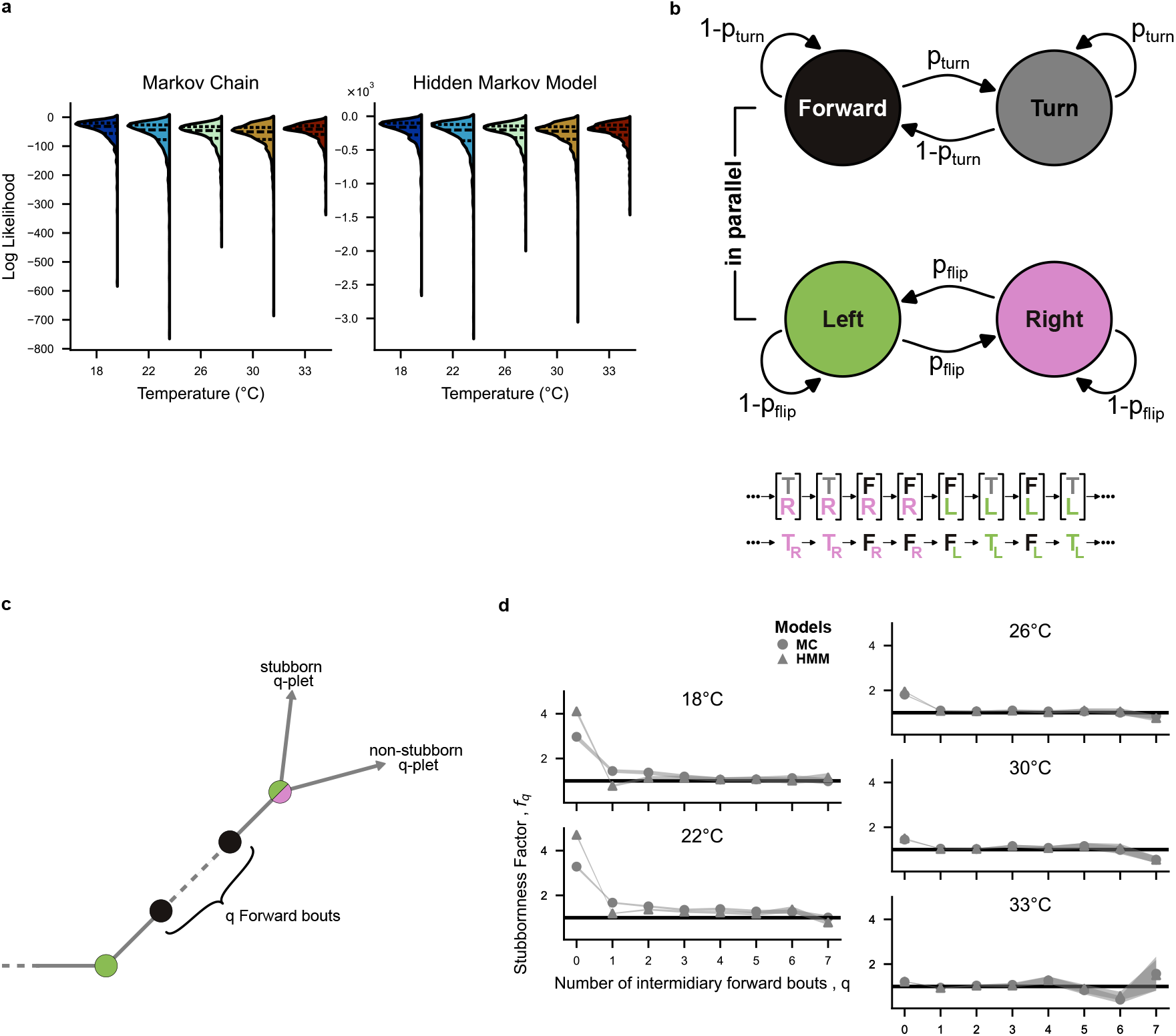
Supplementary panels to Fig.2 - Markovianity: **(a)** Distribution of log Likelihoods (LLHs) for both the Markov chains inferred from thresholded data (left) and Hidden Markov Model (right). For each model and each temperature, LLHs were computed for 100 models inferred from 50% of the trajectories (randomly constructed training set) and on the remaining 50% of the trajectories (testing set). Dashed lines show the quartiles of each distribution. **(b)** Diagram of the 4-state Markov chain used in previous publications [20, 22]. Two Markov Chains run in parallel, with the first chain controlling bout type (forward or turn) and the second controlling direction (left or right). With this model, the system can be in one of four states: [*T, L*], [*T, R*], [*F, L*], [*F, R*], thus left and right states represent internal directional states (not only observed behavioral orientations). **(c)** Diagram illustrating the definition of the stubbornness. For a q-plet of bouts *T*_1_→ *F*→ … → *F* →*T*_2_ with *q* intermediary forward bouts, a stubborn sequence is defined as one where directionality is conserved (i.e. *T* 1 = *T* 2), whilst a non-stubborn sequence will lose the memory of the initial turn (i.e. *T* ≠ 1 *T* 2). **(d)** Evolution of the stubbornness factor *f*_*q*_ (see Eq. 3) with the number of intermediary forward bouts *q*, comparing the Markov Chain inferred from thresholded trajectories (MC, dots) and the Hidden Markov Model (HMM, triangles) trained directly from reorientation angles, for each temperature. The width of the shaded curves represent the estimated error in *stubbornness* factor (see Materials and Methods IV D).

**FIG. S5.**
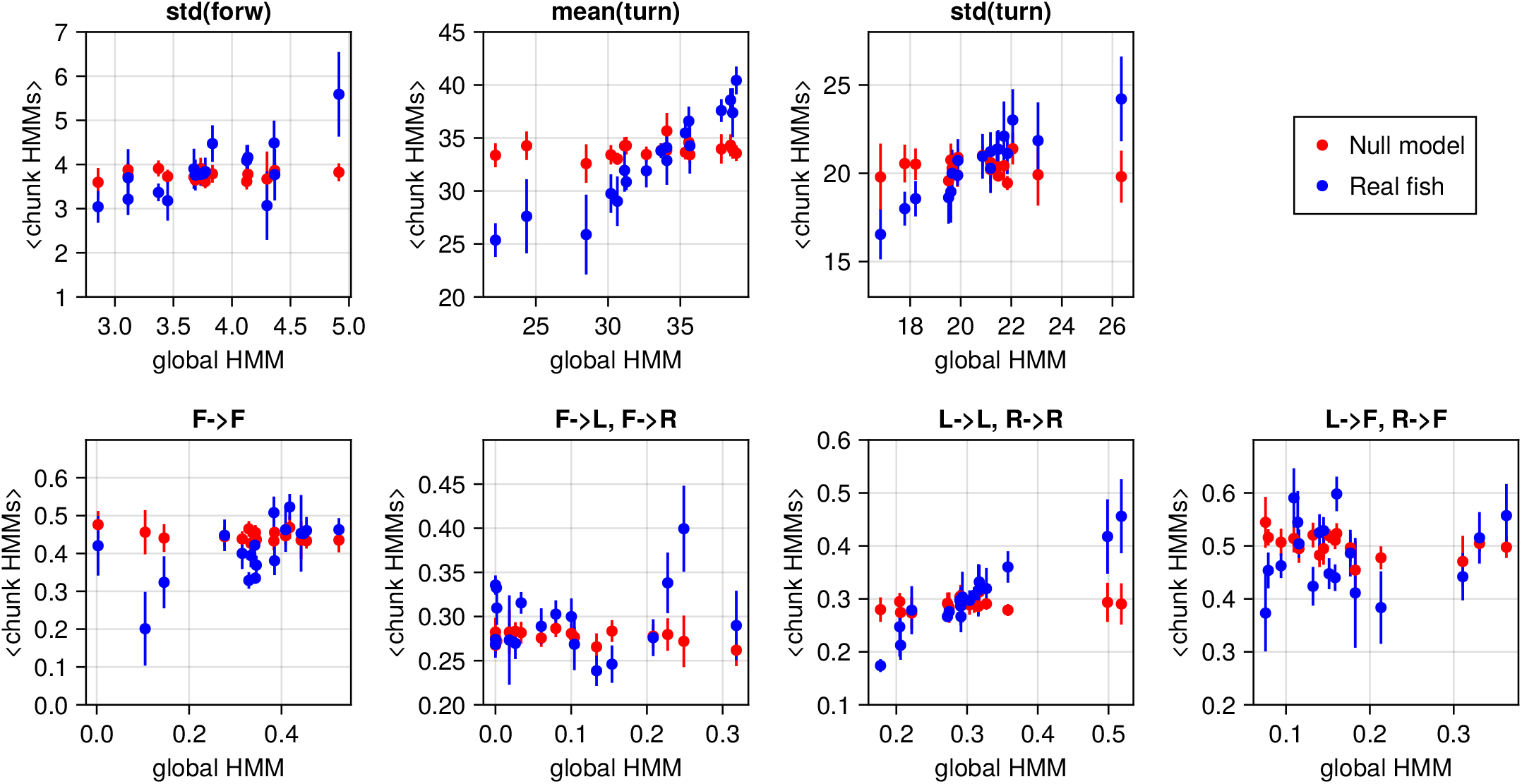
Supplementary panels to Fig.3. Hidden Markov Model parameters inferred from all trajectories from an individual fish, compared with the average parameters inferred from chunks of that fish’s trajectories. All HMM parameters are shown. Each dot represents a fish, with error bars corresponding to standard error of the mean. Blue color corresponds to real individual fish data. Red points are obtained by sampling long trajectories from a single HMM trained on all fish bundled together, thus representing a null model for the fish individuality.

**FIG. S6.**
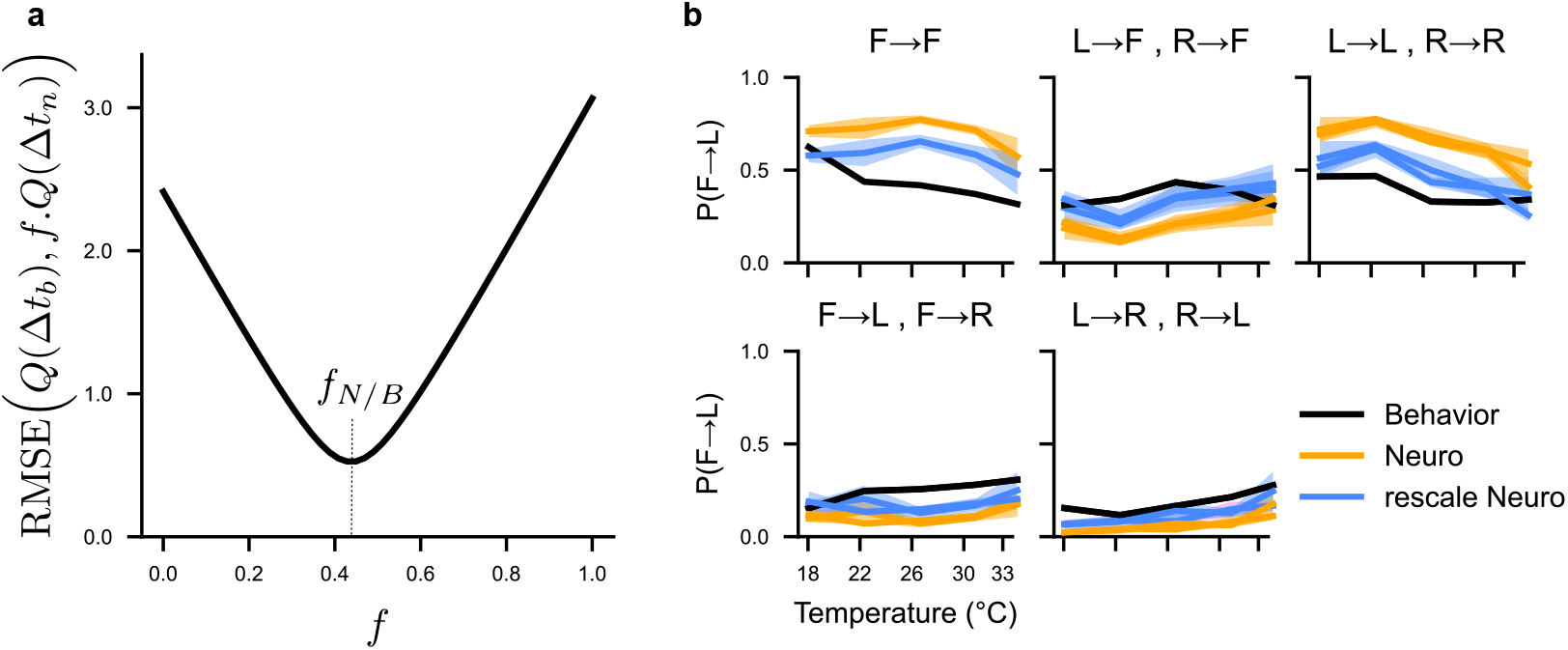
Supplementary panels to Fig.5. **(a)** Root Mean Squared Error (RMSE) between quantiles of the behavior and neuronal sojourn distributions presented in Figure 5a at different values of the rescaling factor *f*. The optimal rescaling factor corresponds the minimal RMSE at *f* = *f*_*N/B*_ ≈ 0.44. **(b)** Comparison of the transition probabilities *P* (*s* → *s*′) between hidden states *F, L*, and *R*, for the behavioral HMM (black), neuronal HMMs (orange), and neuronal HMM rescaled by *f*_*N/B*_ = 0.44 (magenta) at all 5 recorded temperatures. Shaded curves represent the standard error of the mean for all recorded fish at each temperature.

**FIG. S7.**
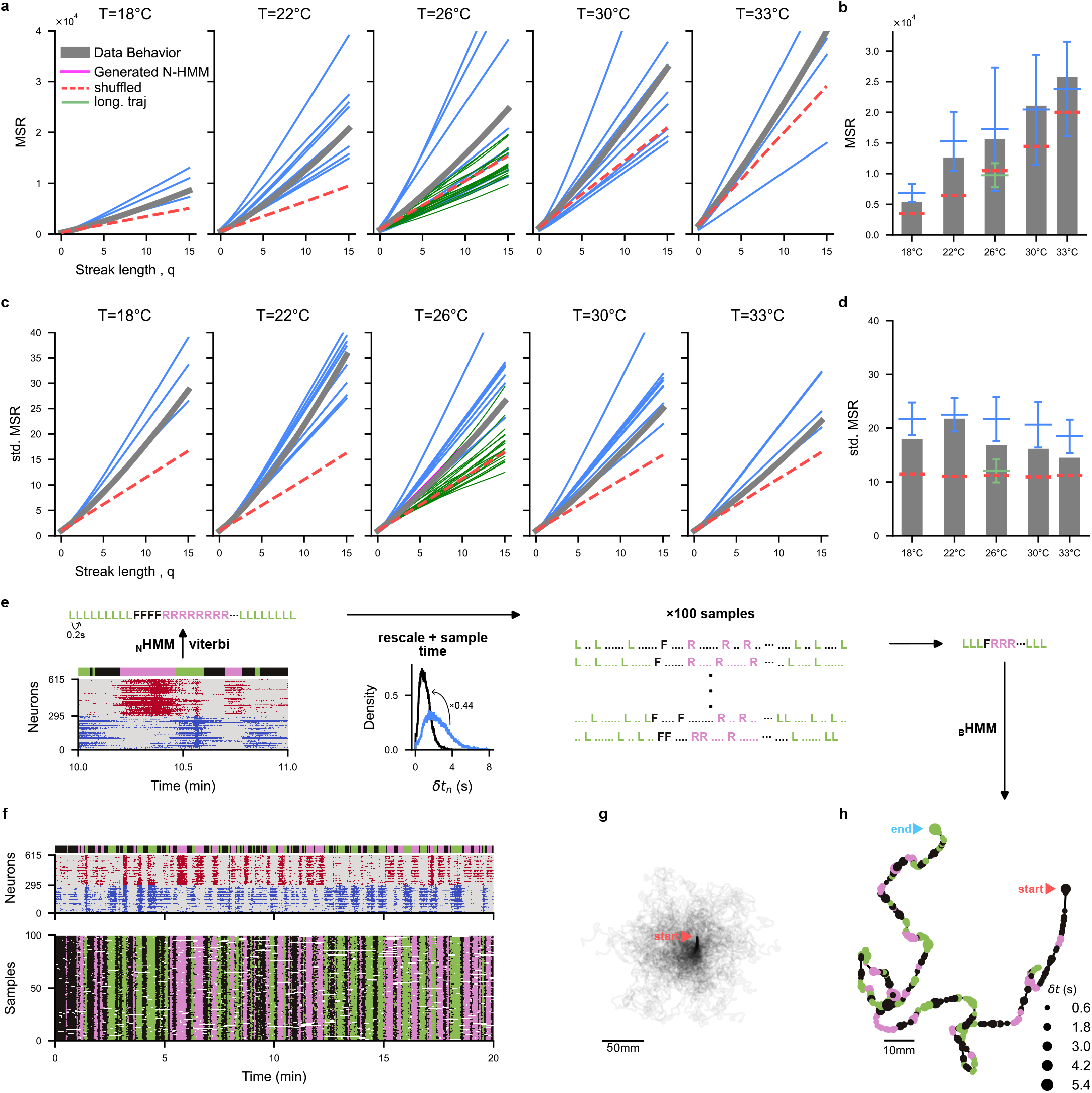
Supplementary panels to Fig.6. **(a)** Mean Square Reorientation (MSR) after *q* bouts from aggregated multiple-fish trajectories at 18-33 ^*o*^C (grey), long-individual trajectories at 26^*o*^ C (green) and generated trajectories from Neural HMM (N-HMM) (blue). Red dashed lines are MSR obtained from shuffled aggregated multiple-fish trajectories. **(b)** Bar plots for the MSR(q=10) for data and N-HMM generated trajectories, with mean (horizontal bars) and standard deviations (vertical bars). **(c-d)** Same as panels a-b but plotting the standardized MSR where trajectories are normalized such that the bout angles have unit variance. See Eq. (A.28). **(e)** Diagram explaining the conversion from neuronal activity to swim trajectory. ARTR activity is first converted into a sequence of forward, left, right hidden states using the Viterbi algorithm on the N-HMM. Time is then re-scaled using the scaling factor identified in Fig 5, and bout sequences are sampled based on the interbout interval distribution. A swim trajectory is constructed for each bout sequence by sampling the bout distances *d*_*n*_ and inter-bout intervals *δt*_*n*_ emission distributions in the behavioral HMM. **(f)** Example recorded ARTR activity at 26^*o*^C (top) and corresponding state sequences after temporal re-scaling and bout sampling (bottom). **(g)** Reconstructed trajectories for each sampled state sequence in panel f. **(h)** Example reconstructed trajectory from the ARTR activity in panel f.

## Notes

https://github.com/ZebrafishHMM2023/ZebrafishHMM2023_CodeAndData/tree/bioRxiv

https://github.com/ZebrafishHMM2023/ZebrafishHMM2023.jl/tree/bioRxiv

### Competing Interest Statement

The authors have declared no competing interest.

